# Single-Molecule Imaging Reveals DNA Shape Read-out by the INO80 Chromatin Remodeler

**DOI:** 10.64898/2025.12.11.693632

**Authors:** Garp Linder, Daniela Horky, Sarah Zernia, Johannes Stigler

## Abstract

ATP-dependent chromatin remodeling enzymes act together to construct complex chromatin architectures around functionally important regulatory regions like promoters and origins of replication. The INO80 complex accurately positions nucleosomes bordering these regions, which is crucial for proper transcription start site selection and efficient replication. How INO80 localizes these nucleosomes and regulates its remodeling activity to produce specific positions mechanistically is unclear, although recent findings suggest a role for DNA shape features and barrier factors. Here, we use single-molecule DNA curtains to directly observe interactions of INO80 with DNA, nucleosomes and barrier factor Reb1. We show that INO80 recognizes DNA shape and strongly binds to shape features that are enriched in promoters and ORIs. Moreover, we find that INO80 performs 1D searches and intersegmental transfers to dynamically interact with nucleosomes and Reb1, but cannot bypass them, indicating that they could confine INO80 within promoters and thereby increase engagement with flanking nucleosomes. Our findings reveal how INO80’s target-site search and remodeling activity are influenced by DNA shape recognition and molecular architecture and elucidate how remodelers can integrate different kinds of information to transform the chromatin landscape.

## INTRODUCTION

Manipulation of nucleosomes, the core unit of chromatin, constitutes a major share of eukaryotic genome regulation and results in the formation of diverse chromatin states [1–3]. While histone modifications are modulated by different epigenetic modulators, the positioning and histone composition of nucleosomes is mainly directed by four families of ATP-driven chromatin remodeling complexes: SWI/SNF, ISWI, CHD, and INO80 [4]. Significant research efforts conducted in the last two decades in *S. cerevisiae* have revealed that these protein families work in concert to generate specific chromatin architectures around important regulatory regions like promoters and origins of replication (ORI) [5–7]. The architectures produced are characterized by a nucleosome depleted region (NDR) flanked by two well-positioned nucleosomes, which in context of promoters were coined the +1 (over the transcription start site) and *−*1 (opposite side) nucleosome. Nucleosomes further up- and downstream show regular spacing and are phased relative to the +1*/ −* 1 nucleosomes [3, 8]. Maintenance of this architecture seems to be critical, as disruption has been shown to promote cryptic transcription and reduce replication efficiency [7, 9]. The only chromatin remodeler of the four families to accurately reproduce *in-vivo*-like +1*/ −* 1 nucleosome positions in a *in-vitro* positioning assay was INO80 [10], a remodeler with nucleosome and hexasome sliding activity and proposed functions in multiple other pathways, such as replication and DNA repair [11–13]. INO80 is present almost exclusively on nucleosomes flanking NDRs [14] and its removal enhances cryptic transcription [5, 9, 15], making it a prime candidate for genome wide positioning of +1*/ −* 1 nucleosomes. Results from selective and simultaneous depletion of different remodelers *in vivo* support this notion [5, 16].

In *S. cerevisiae*, INO80 is comprised of 15 subunits that can be grouped into three structural modules, hereinafter referred to as the Core-, A-, and N-module [17] (fig. 1A). All three modules are connected by the Ino80 subunit, which also contains the Snf2-like ATPase motor. During repositioning of nucleosomes, INO80 forms an extended configuration, where the Core-module binds to the nucleosome, while the A- and N-modules sit in front of the ATPase motor and interact with the extranucleosomal DNA, which is translocated towards the motor [18, 19]. Using this architecture INO80 positions nucleosomes by pulling them into the NDR [5].

**FIG. 1.**
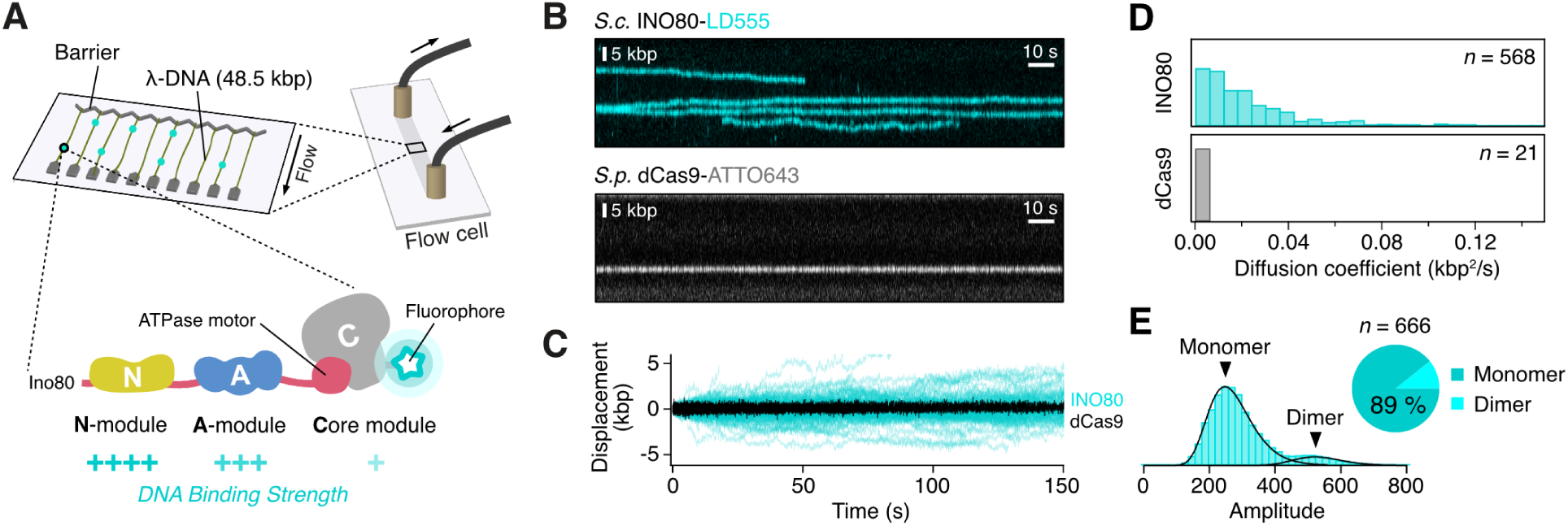
Single-molecule imaging of INO80 diffusion. **(A)** Overview of DNA curtains setup and INO80 submodule organization. [18] **(B)** Kymograms of INO80 labeled with LD555 diffusing along DNA (top) and dCas9 labeled with Atto643 binding statically to DNA (bottom). **(C)** Aligned trajectories of all analyzed INO80 (cyan) and dCas9 (black) translocation events. **(D)** Histogram of diffusion coefficients for INO80 and dCas9. **(E)** Left: Histogram of measured amplitudes for all localized particles, showing two fitted peaks for monomers and dimers. Right: Relative fraction of monomeric and dimeric localizations.

The ability of INO80 to accurately slide nucleosomes to certain positions necessarily implies negative regulation of the sliding process through external cues. INO80 has been shown to position and phase nucleosomes relative to other proteins bound to DNA, e.g., general regulatory factors (GRF) Reb1 and Abf1 at promoters and ORC at ORIs, most likely through a ruler-like readout [7, 20]. How this is facilitated mechanistically is unknown. Interestingly, INO80 is also able to position +1*/ −* 1 nucleosomes in the absence of such ‘barrier proteins’, solely based on DNA sequence inputs. Analysis of final +1 nucleosome positions remodeled *in vitro* by INO80 revealed distinct DNA shape profiles, which changed upon mutation of different subunits [10]. Another study observed that INO80 remodels nucleosomes with rigid linkers more slowly [21]. This suggests that INO80 can regulate its remodeling activity through recognition of DNA shape features. DNA shape sensing of proteins is mostly facilitated by interactions with the phosphate backbone of the DNA and reads out characteristics of the DNAs three-dimensional structure like minor groove width, propeller twist or DNA curvature [22]. Although structural and biochemical data has shown that INO80 can recognize these structures [10, 19], it is not known how they influence INO80 binding dynamics.

Additionally, while the target-site search of transcription factors (TF) and other small DNA-binding proteins (DBP) has been studied for decades [23–25], a comprehensive model for how chromatin remodelers, including INO80, locate their target nucleosomes in a busy chromatin landscape is still missing. This is in part due to the limitations of classical biochemical bulk assays, which are limited in their ability to observe interaction dynamics while bound to the substrate and therefore cannot resolve molecular details of target site search. Recent single-molecule studies of RSC, ISW2 and SWR1 *in vitro* have presented first insights by showing that these remodelers, similar to most DNA-binding proteins, use 1D diffusion to locate target proteins like nucleosomes on DNA [26, 27]. Similar *in-vitro* studies are still missing for INO80.

In this study, we used single-molecule DNA curtains to address INO80’s target site search, DNA binding preferences and interaction kinetics with Reb1 and H2A.Z-containing nucleosomes. Additionally, we performed sliding assays to reveal the effect of DNA shape recognition on nucleosome remodeling. We demonstrate that INO80 uses 1D diffusion by helical sliding to move along DNA and can jump between DNAs using intersegmental transfers. We observe highly DNA shape dependent binding and diffusion kinetics with stronger binding and slower diffusion speeds on DNA sequences with high propeller twist angles, which are frequent in promoter NDRs [28]. Using truncation mutants we identify the A-module as a sensor for these sequences and show that their recognition within nucleosomal linker DNA leads to new intermediates during remodeling. Additionally, we present dynamic interactions of INO80 with nucleosomes and Reb1 and find that both act as barriers to INO80 diffusion, indicating that they could trap INO80 in NDRs and increase engagement with the flanking +1*/ −* 1 nucleosomes. Together, this study reveals detailed insights into how DNA shape, molecular composition and interaction partners govern INO80’s kinetics on DNA, laying the foundation for a comprehensive target site search model and providing a framework for DNA shape dependent nucleosome positioning.

## RESULTS

### INO80 diffuses along DNA

We expressed and purified the *S. cerevisiae* multi-subunit INO80 complex endogenously and labeled it fluorescently at the C-terminal ybbR-tag of the Ino80-subunit (fig. 1A). To investigate INO80’s DNA binding and translocation, we used single-molecule DNA curtains, which allow the visualization of individual INO80 complexes on DNA in real time and with high throughput. We used flow cells with nano-fabricated chrome barriers that were passivated with a lipid-bilayer, and assembled a double-tethered DNA curtain using end-modified *λ*-DNA molecules (48.5 kbp), as previously described [29] (fig. 1A). A microfluidic system in combination with total internal reflection (TIRF) microscopy allowed us to apply fluorescently labeled INO80 complexes to the flow cell and record association and dissociation events of individual molecules.

INO80 bound stably to *λ*-DNA with a mean lifetime of 113 *±* 9 s and slowly moved along the DNA in a non-directed fashion resembling a random walk (fig. 1B,C). As a control, we applied a catalytically dead Cas9 mutant (dCas9) to the DNA, which, as expected, bound to its target sites and showed no movement along the DNA (fig. 1B,C, fig. S1A). The analysis of single-molecule diffusion coefficients revealed a wide distribution for INO80 with a median diffusion coefficient over 30 times higher than for dCas9 (0.017 *±* 0.001 kbp^2^*/*s for INO80 vs. (5.5 *±* 3.8) *×* 10^−4^ kbp^2^*/*s for dCas9, fig. 1D). We hence conclude that INO80 translocates along DNA by 1D diffusion.

A small subset of complexes diffusing on DNA showed noticeably higher fluorescence intensities, and photo bleaching analysis revealed that these complexes bleached in two steps (fig. S1B). Analysis of all localized intensities displayed two distributions, indicating that at least *≈*11 % of DNA-bound INO80 complexes were dimers (fig. 1E). However, the diffusion coefficients and lifetimes of dimeric complexes showed no significant differences compared to monomers (fig. S1C), implying that these complexes are functionally indistinguishable from monomers in respect to their DNA binding.

### INO80 slides along DNA using a rotationally coupled mechanism

Having established that INO80 uses 1D diffusion, we sought to uncover how INO80 moves along DNA mechanistically. DBPs have been shown to slide, hop or use a mix of both mechanisms [27, 30]. This profoundly impacts the way the proteins interact with obstacles and interaction partners along the DNA and is therefore crucial for understanding their target site search mechanism [31]. To test whether INO80 hops or slides we compared INO80 diffusion at 120 mM KGlu with diffusion at 40 mM KGlu. As hopping proteins have to dissociate and re-associate continuously while moving, their diffusion coefficient is positively correlated to their off-rate and hence salt-dependent [30]. As expected, with reduced salt concentration, INO80’s median lifetime increased four-fold from 15.1 s to 61.9 s. Its median diffusion coefficient decreased slightly, but the change was not significant (0.012 *±* 0.002 kbp^2^*/*s to 0.017 *±* 0.001 kbp^2^*/*s; *p* = 0.055, Mann-Whitney U test, fig. 2A). This implies that INO80 movement is mostly salt-independent and based on sliding.

**FIG. 2.**
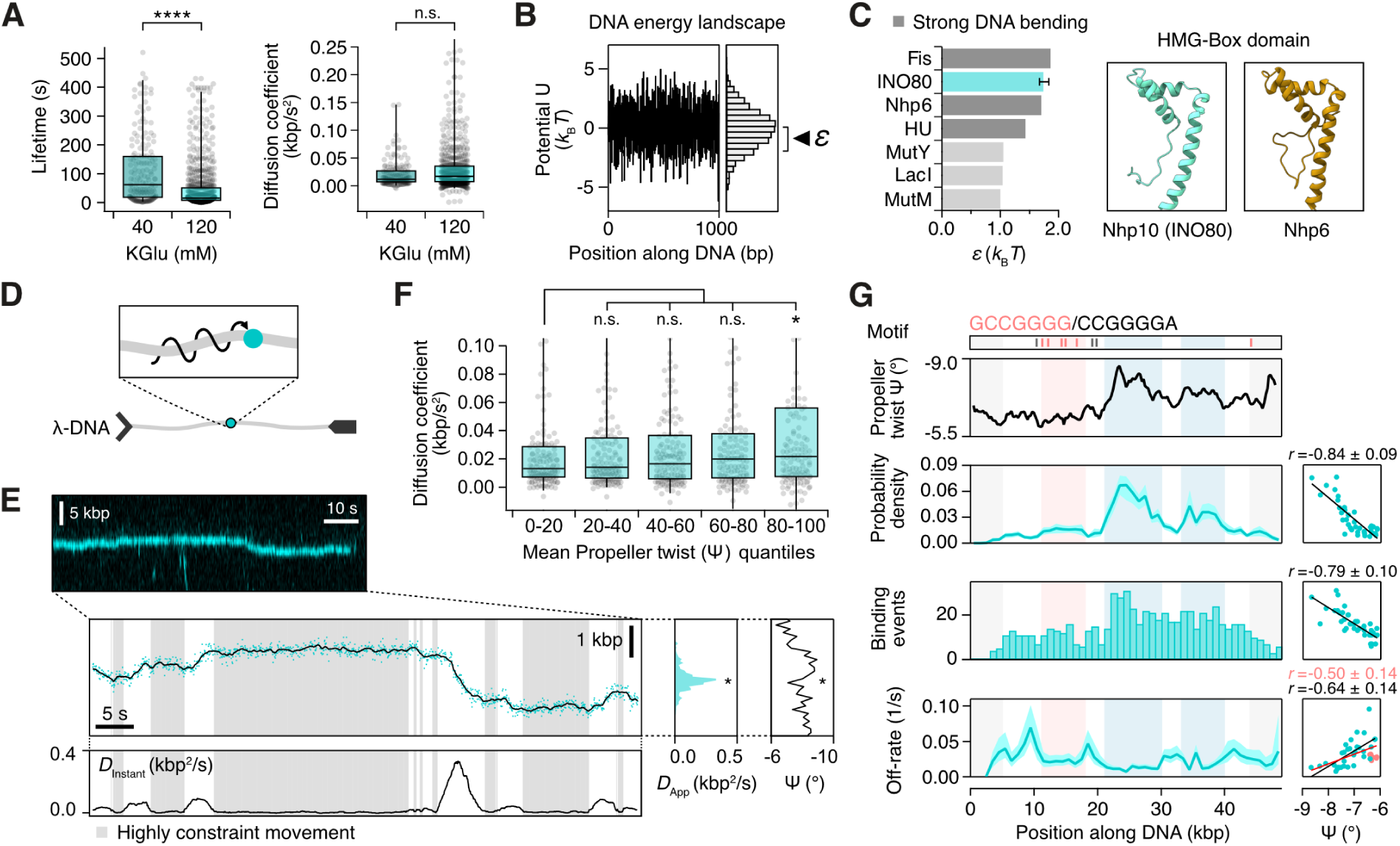
INO80 slides along the helical axis and strongly binds to DNA with high negative propeller twist Ψ. **(A)** Comparison of lifetimes and diffusion coefficients of INO80 on *λ*-DNA for different salt concentrations. Significance was determined using a Mann-Whitney U test. *n*_40KGlu_ = 167. *n*_120KGlu_ = 677 **(B)** Example energy langscape felt by particle with free-energy roughness *ε*. **(C)** Comparison of free-energy roughness for different DBPs moving along the helical pitch of the DNA, obtained from [32, 35]. Dark grey shading indicates proteins that strongly bend DNA. Right: Predicted Alphafold structure of Nhp10 (AF-Q03435-F1-v6) and NMR structure of HMG-Box of Nhp6 (PDB:1LWM). **(D)** Model for INO80 diffusion. INO80 slides along the helical pitch of the DNA. **(E)** Top: Kymogram showing a diffusing INO80 complex exhibiting variable constraint depending on position along the DNA. Bottom: Respective trajectory time trace with instantaneous diffusion coefficient (window size: 5 s). Shaded areas indicate highly constraint movement (*D*_Instant_ *<* 0.01 kbp^2^*/*s), dots show localizations, black line shows positions determined using a Kalmanfilter. Right: Apparent diffusion coefficient of the respective molecule and average propeller twist of the DNA depending on position. **(F)** Diffusing INO80 complexes were sorted into quintiles depending on the mean propeller twist (Ψ) they encountered during diffusion along *λ*-DNA. Significance was determined using a Mann-Whitney U test. *n* = 665. **(G)** Nhp10-Motifs, average propeller twist (1 kbp sliding window) and different binding parameters along *λ*-DNA. Shaded areas: close to chrome barriers with unreliable data (grey), accumulation of Nhp10 motifs (red), high negative propeller twist (blue). Grey shaded areas were not included in correlation analysis. *n* = 677. Right boxes: correlation of binding parameters with average propeller twist and Pearson’s correlation coefficient.

Sliding proteins can diffuse by moving along the helical axis in a rotational fashion [32] or by moving linearly over the DNA employing a rotationally decoupled mechanism [33]. To get insights into INO80’s diffusion mode we analyzed the free-energy landscape along the protein’s path of diffusion by calculating its roughness *ε*, which causes retardation of the effective diffusion speed due to protein interactions with the DNA (fig. 2B). Using the median diffusion coefficient measured at 120 mM KGlu and a radius of *≈*100 Å for INO80 (estimated from published cryo-EM structures), we inferred a free-energy roughness of *≈*1.75 *k*_B_*T* for rotationally coupled diffusion, while linear diffusion would predict a much higher roughness of *≈*3.15 *k*_B_*T*. Consistent with rotational movement but incompatible with linear movement, models predict roughness values between 0.5 to 2 *k*_B_*T* for efficient target site search [34], which has been confirmed for a wide variety of DBPs [32, 35]. Additionally, the value for rotationally coupled movement closely fits the free-energy roughness determined for Nhp6, a HMGB-type protein closely related to the DNA binding subunit Nhp10 within the INO80 N-module [36] (fig. 2C). Nhp6 bears a similar HMGB-type box domain as Nhp10, has been shown to slide rotationally and induces high bending angles into the DNA during binding [35]. Hence, we conclude that also INO80 moves by rotationally-coupled sliding along DNA and possibly bends the DNA in the process (fig. 2D).

### INO80 transfers between DNA strands

On the DNA curtains chip, DNA strands are juxtaposed, but tethering is not always perfectly parallel and multiple DNAs can tether within the same well (fig. S2A). This, and naturally occurring thermal fluctuations allow the *λ*-DNA molecules to come into close contact and enables proteins, if able, to jump between DNA strands (jumps in *trans*). We frequently observed these jumps by INO80, which were characterized by an immediate change in *x*-position (fig. S2B). We used a Hidden-Markov-Model (HMM) to estimate jump timepoints and compared displacements around these timepoints with random displacements of INO80 complexes diffusing along DNA (fig. S2C,D). Unlike random displacements, which were normally distributed, *x*-displacements at jump timepoints showed a bimodal distribution. Intriguingly, the corresponding *y*-displacements showed no significant difference compared to random y-displacements. This indicates that for these kinds of jumps the complex does not undergo complete dissociation, 3D diffusion and re-association during the transfer, but moves directly from one to the other DNA strand during random DNA-DNA collisions, thus mechanistically fulfilling the definition for inter-segmental transfers.

Moreover, we also observed jumps between distant positions on the same DNA strand (*cis*), visible by changes exclusively in *y*-position (fig. S2E). The DNA is stretched to *≈*74 % of its contour length along the *y*-direction. Reptations and looping are hence highly limited in this direction and far distant regions of the same DNA strand cannot come into close proximity. To perform *cis* jumps, INO80 must therefore dissociate, diffuse in 3D space and re-associate at a different position on the same DNA. These *cis* jumps should not be confused with the aforementioned systematic hopping diffusion mechanism, but rather represent transitions from 1D to 3D to 1D, where the protein by chance reassociates with the same DNA and can therefore cover much larger distances.

### INO80 accumulates on DNA with high negative propeller twist

During closer analysis of INO80 trajectories, we noticed that diffusing INO80 complexes showed varying mobility on *λ*-DNA over time. This non-random movement was represented by transitions between highly diffusive and highly constrained segments within 1D trajectories of single complexes (fig. 2E), an unexplained behavior also observed for RSC and ISW2 [27]. Given that INO80 is hypothesized to recognize DNA shape features and that our setup allowed us to analyze the DNA sequence background of diffusing complexes, we tested whether INO80 binding dynamics are influenced by DNA shape properties. We chose propeller twist as it was singled out as one of the most relevant DNA shape parameter to explain nucleosome placement by INO80 in a previous study [10]. The propeller twist Ψ measures the negative angle between the two bases of a base-pair and is correlated with the electrostatic potential (EP) of the minor groove, which is emerging as a major determinant of DNA shape read-out [37] (fig. S2F). We sorted diffusion coefficients of individual complexes into quintiles depending on the average propeller twist that they had encountered along their trajectory and compared them (fig. 2F). The analysis revealed a clear trend towards faster diffusion at low Ψ with the lowest and highest quintiles showing a significant difference. Next, we determined the localization probability density, the initial binding position and the off-rate of INO80 complexes dependent on the DNA position and correlated it with the respective Ψ at this position (fig. 2G). Strikingly, INO80 was strongly enriched on DNA sequences with high Ψ, showed more initial binding to and longer lifetimes in these DNA regions. Given the strong correlations we wondered whether other DNA shape features could explain INO80-DNA kinetics even better. We tested 11 additional established intra-base pair and inter-base pair features, as well as two minor groove parameters predicted by Deep DNAshape [38]. No features outperformed propeller twist, but EP, stretch, opening, slide and rise performed equally well (fig. S2G). Together, this supports INO80’s ability to directly recognize DNA shape features, and, additionally, that its binding kinetics are dominated by these features.

INO80 also showed lower off-rates in a segment of the *λ*-DNA between 12 to 18 kbp, which exhibits a comparably low Ψ but contains multiple binding motifs found for INO80-subunit Nhp10 [39] (fig. 2G, red shaded area). This suggests that the N-module might additionally weakly recognize sequence motifs during 1D diffusion, which may be relevant for different cellular functions of the complex besides remodeling.

### Removing INO80’s N-Module speeds up kinetics but retains DNA shape dependence

Having revealed that INO80 displays highly DNA shape dependent binding kinetics, we wondered how DNA shape recognition by INO80 is achieved. Given that the N-module is highly species dependent and that its removal keeps INO80’s ability to reposition nulceosomes largely unaffected [10, 19], we hypothesized that a mutant, missing the N-module and the N-terminal end of the Ino80 subunit would still retain its DNA shape readout capabilities. We chose an established ΔN-INO80 mutant from *C. thermophilum* that has been shown to be highly conserved in structure to *S. cerevisiae* INO80 and which produces similar +1 nucleosome positions [10, 19] and applied it to DNA curtains. This mutant will be referred to from here on as *Ct.* ΔN-INO80. If no species identifier is given, the respective variant mentioned is a *S. cerevisiae* protein or mutant.

Similar to the *Sc.* wildtype, *Ct.* ΔN-INO80 readily bound to *λ*-DNA and moved along the DNA (fig. 3A), but displayed much faster diffusion dynamics with a median diffusion coefficient over 10 times higher than the *Sc.* wildtype complex (0.200 *±* 0.009 kbp^2^*/*s vs. 0.017 *±* 0.001 kbp^2^*/*s; fig. 3B). Compared to the *Sc.* wildtype the calculated free-energy roughness for *Ct.* ΔN-INO80 was halved (0.86 *±* 0.15 *k*_B_*T* vs. 1.84 *±* 0.07 *k*_B_*T*) and consistent with values for DBPs without strong DNA bending (fig. S3A). This validates our earlier observations that diffusion speed in the wild type complex is mainly dominated by the high free-energy roughness of Nhp10’s non-sequence-specific HMG-boxes. Remarkably, the diffusion speed of *Ct.* ΔN-INO80 complexes also showed a stronger correlation with the mean propeller twist angles of the encountered DNA than the *Sc.* wildtype complex (fig. S3B). Analysis of single trajectories revealed that steep transitions from high to low Ψ on the DNA acted as strong barriers for *Ct.* ΔN-INO80, which ‘trapped’ the protein in DNA regions with high Ψ (fig. 3C). Similar to the *Sc.* wildtype complex, other binding parameters of *Ct.* ΔN-INO80 also correlated strongly with Ψ (fig. S3C), indicating that recognition of DNA shape is independent of the *Ct.* N-module and most likely an evolutionary conserved property of the combined Core- and A-module. As expected, the subset of long-living complexes near Nhp10 binding motifs also disappeared (fig. S3C).

**FIG. 3.**
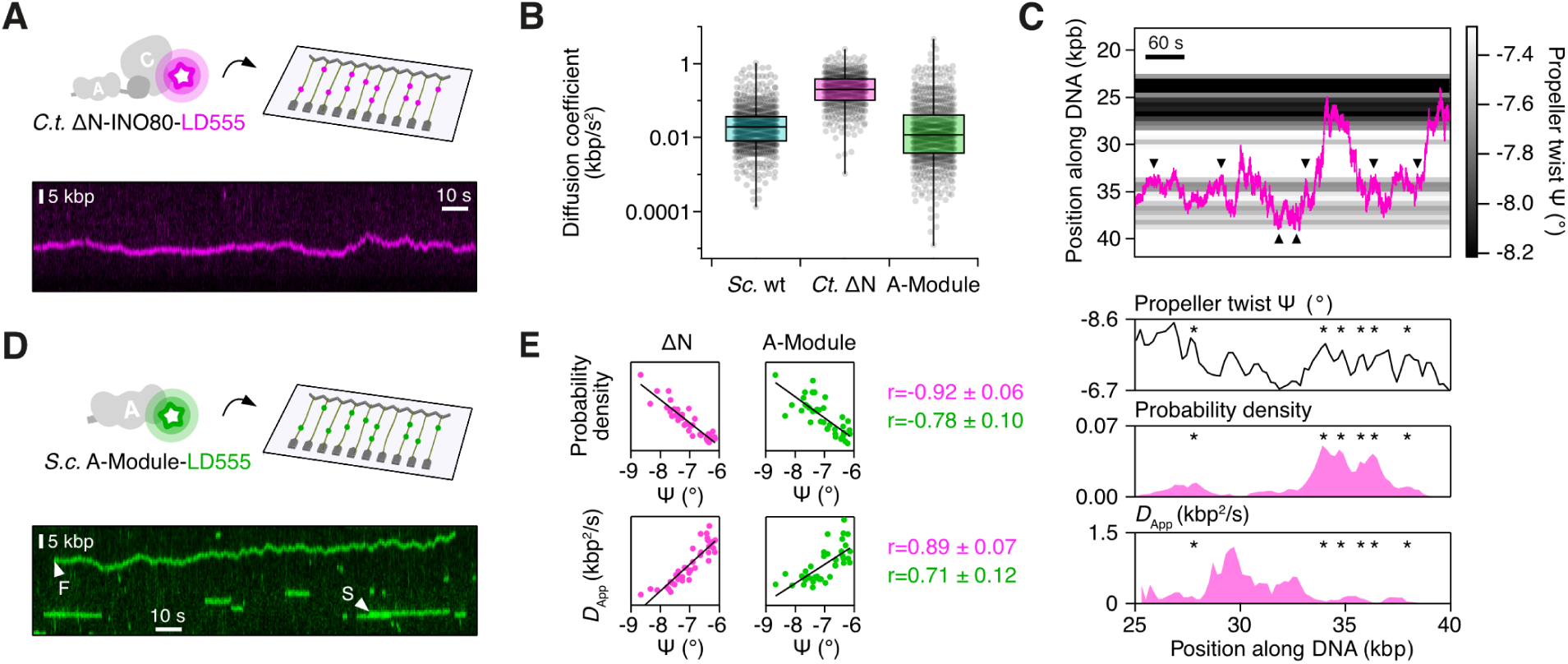
The A-module is a DNA shape sensor. **(A)** Top: Schematic of *Ct.* ΔN-INO80 and DNA curtains assay. Bottom: Kymogram of LD555-labeled *Ct.* ΔN-INO80 diffusing along DNA. **(B)** Diffusion coefficients of all INO80 variants. **(C)** Top: Single trajectory of *Ct.* ΔN-INO80 complex. Arrows indicate where the complex is blocked at transitions from high to low propeller twist. Bottom: Position dependent propeller twist of *λ*-DNA, probability density and *D*_App_ for trajectory shown above. Asterisks indicate DNA positions with high propeller twist occupied by the complex. **(D)** Top: Schematic of INO80 A-module and DNA curtains assay. Bottom: Kymogram of LD555-labeled A-module diffusing slowly (S) or fast (F) along DNA. **(E)** Correlation of probability density of localizations and apparent diffusion coefficient with average propeller twist and Pearson’s correlation coefficient for *Ct.* ΔN-INO80 (magenta, *n* = 1180) and INO80 A-module (green, *n* = 851).

We again observed transfers between DNA strands without a change in *y*-position similar to those of the *Sc.* wildtype, suggesting that the *Ct.* ΔN-INO80 complex architecture still enables intersegmental transfers (fig. S3D).

### The A-module is a sensor for DNA shape

After establishing that the N-module was redundant for DNA shape sensing, we turned our attention to the conserved A-module. The position of the A-module directly in front of the ATPase motor and their shared connection via the Ino80 HSA/post-HSA domain suggest that it might play a crucial role in recognizing DNA shape and regulating nucleosome sliding activity allosterically [19]. We therefore purified and labeled the *S. cerevisiae* A-module, using a truncated version of Ino80 (440–598), which binds all relevant subunits and forms a structurally fully cohesive A-module [19].

A-module complexes on DNA curtains bound strongly to *λ*-DNA and showed very diverse kinetics (fig. 3D). The majority of the complexes were almost static (S), while some complexes diffused very rapidly (F). Surprisingly, on average, A-module complexes diffused slightly slower than the *Sc.* wildtype complex and much slower than *Ct.* ΔN-INO80 (fig. 3B) with a median diffusion coefficient of (9.7 *±* 0.8) *×* 10^−3^ kbp^2^*/*s. Nevertheless, analysis of apparent diffusion speeds at specific positions revealed that kinetics were again dependent on DNA shape, with slow diffusion at positions with high Ψ (fig. 3E, bottom; fig. S3E). Accordingly, also probability density, initial binding positions and off-rates were highly dependent on the propeller twist of the DNA (fig. 3E, top; fig. S3F). The fact that diffusion kinetics were very heterogeneous, highly DNA-shape dependent, but on average almost static compared to *Ct.* ΔN-INO80, suggests multiple DNA binding configurations, which in the full complex are additionally modulated by the associated Core-module. In line with this, structures of the *Ct.* A-module show, that it can bind to DNA in at least two different states, with or without additional protein-DNA contacts at the N-terminal end of the module [19, 40]. Taken together, our data confirms the A-module as a DNA shape sensor that confers strong binding preferences for DNA with high Ψ to the INO80 complex.

### Influence of DNA-shape recognition on INO80 remodeling

With a clear picture of shape-dependent DNA kinetics by INO80 emerging, we wondered whether DNA shape recognition by the A-module might also influence remodeling kinetics. In gel-based sliding assays using nucleosomes with sufficiently long linker DNA, INO80 translocates end-positioned nucleosomes into the center of the DNA [41]. A previous study had shown reduced remodeling speed in gel-based sliding assays when native DNA sequences with high Ψ were incorporated into linker DNA [21]. However, the random distribution of high Ψ motifs across the native linker sequences made it difficult to determine which interaction is responsible for this effect. During a complete reaction cycle, the DNA is pushed 40 bp along different protein-DNA binding interfaces, including the ATPase motor. Another study showed that sliding is strongly reduced if a high Ψ motif is placed at SHL *−*6/7, where the ATPase motor contacts the DNA [19]. We therefore designed nucleosome constructs with 80 bp linker DNA (0N80), whose average Ψ is uniformly distributed across the whole linker sequence and inserted a 10 bp long poly(dA:dT)-tract, which due to its unique structure exhibits extremely high Ψ (fig. 4A). We chose SHL *−*12/13 as the position for insertion, because only the A-module and N-module will contact it during the remodeling process, according to current models and information from cryo-EM structures and footprinting [19, 40, 42, 43]. Improved separation of nucleosome species using gradient gels allowed us to analyze the emerging remodeling intermediates.

**FIG. 4.**
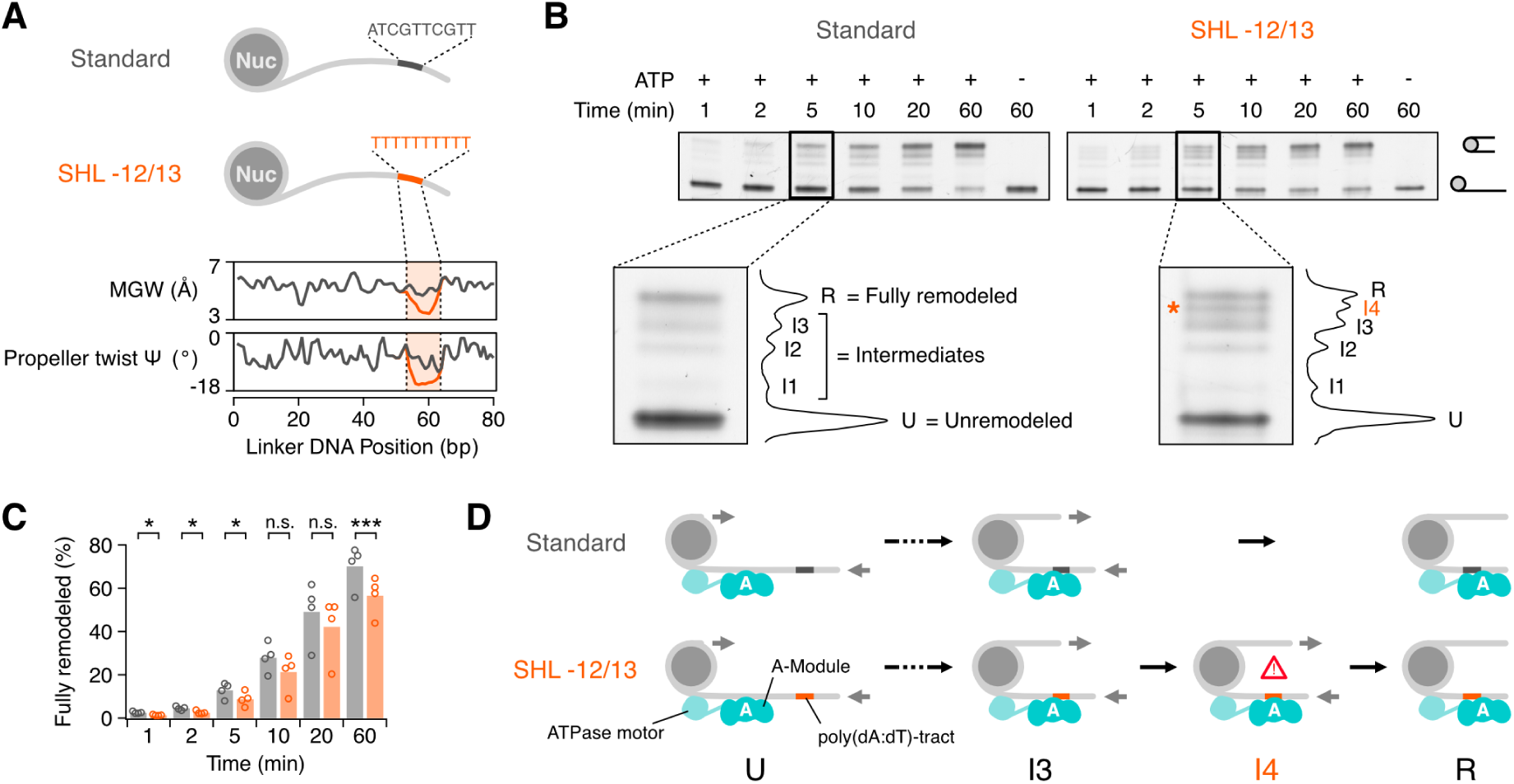
Effect of DNA shape sensing on nucleosome remodeling. **(A)** Top: Schematic of 0N80 nucleosome design for gel-based sliding assays. Sequence exchanged at SHL *−*12/13 is shown. Bottom: Predicted DNA shape parameter for linker DNA of both nucleosome constructs (dark grey: Standard; orange: SHL *−*12/13). **(B)** Top: Native gel showing remodeling of 0N80 nucleosomes over time depending on linker DNA. Bottom: Exemplary band pattern showing nucleosome species originating during remodeling, with associated band intensity plot. Asterisk (*) indicates emergence of an additional remodeling intermediate, when an A-tract is introduced at SHL *−*12/13. **(C)** Comparison of fully remodeled nucleosome fraction for Standard and SHL *−*12/13 nucleosomes. Significance was determined using a two-tailed paired t-test. *n* = 4. **(D)** Schematic of the remodeling reaction showing the placement of A-module, ATPase motor and poly(dA:dT)-tract for different remodeling intermediates, assuming *≈*10 bp remodeling steps [44, 45].

Remodeling of nucleosomes without poly(dA:dT)-tract produced four additional bands representing intermediates *I1* –*I3* and the fully remodeled 40N40 nucleosome *R* (fig. 4B), in line with a translocation step size of *≈*10 to 20 bp previously reported for INO80 [44, 45]. Strikingly, remodeling of the SHL *−*12/13 nucleosome produced an additional intermediate *I4* close to the fully remodeled product in all replicates, indicating an additional pause or termination point of the sliding process between *I3* and *R*. Furthermore, the fraction of fully remodeled product was smaller for all timepoints, when the poly(dA:dT)-tract was present (fig. 4C). Reconstruction of the approximate location of the poly(dA:dT)-tract within *I4* (assuming *≈*10 bp steps) places it in direct contact with the A-module (fig. 4D). Our results therefore suggest that recognition of poly(dA:dT)-tracts through the A-module can lead to abortion of the translocation process and hence inhibition of remodeling.

### INO80 dynamically interacts with nucleosomes

Next, we focused our attention on the role of interaction partners of INO80 within the context of target-site-search and remodeling. *In vivo*, INO80 preferentially interacts with and remodels +1 nucleosomes, which are enriched for the histone variant H2A.Z [46]. We therefore assembled labeled H2A.Z-containing nucleosomes on *λ*-DNA and generated sparsely chromatinized DNA curtains to which we added INO80 (fig. 5A).

**FIG. 5.**
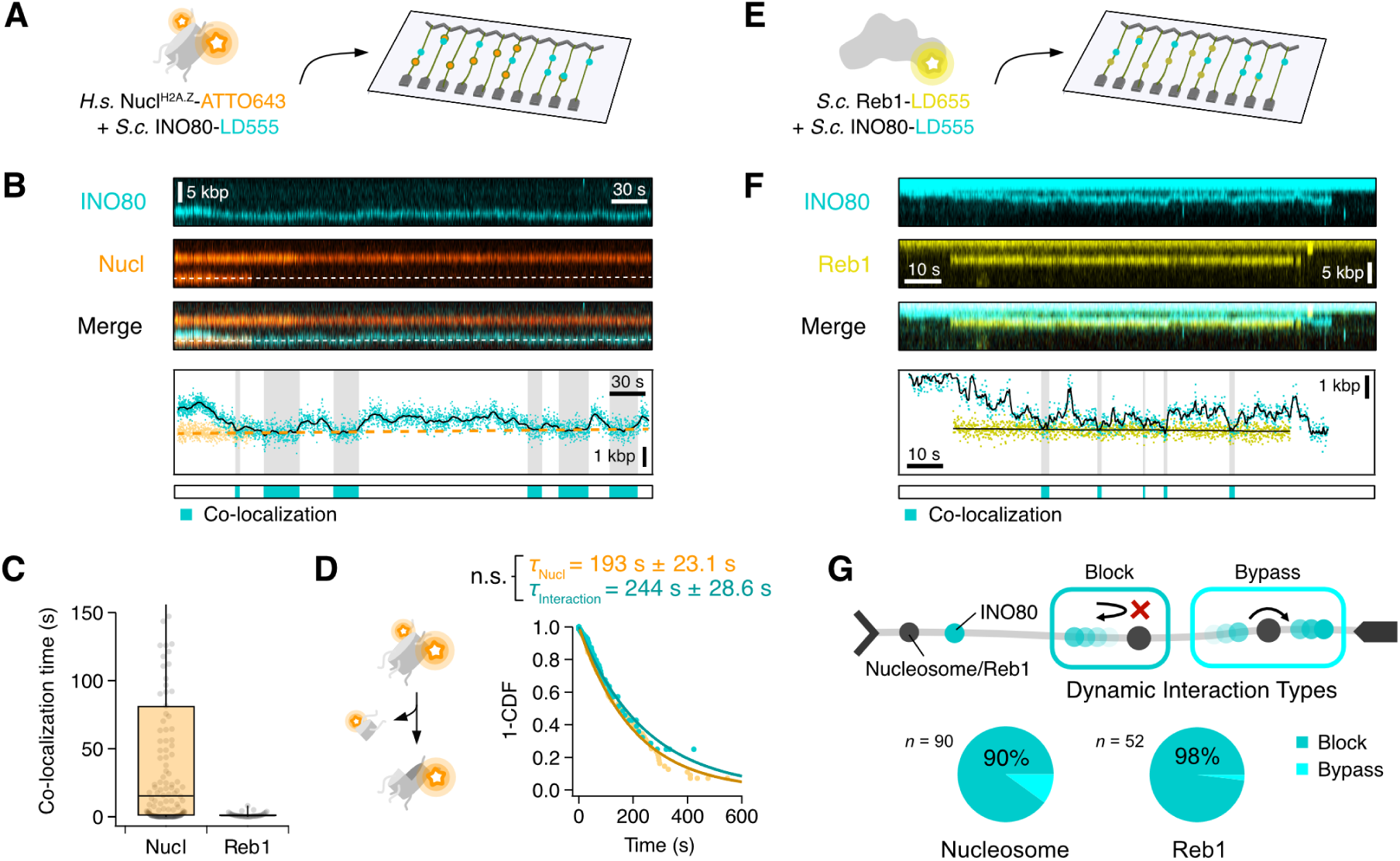
INO80 interactions with nucleosomes and Reb1. **(A)** Schematic of chromatinized DNA curtains assay. **(B)** Top: Kymogram of an INO80 complex (cyan) co-localizing with a nucleosome (orange). Bottom: Corresponding time traces with co-localizations indicated by shaded areas. **(C)** Comparison of INO80 colocalization times with nucleosomes (*n* = 142) and Reb1 (*n* = 34). **(D)** Left: Schematic for eviction assay. Right: Survival plot of nucleosome lifetimes alone or while bound by INO80. Rates were compared with Log-Rank test. **(E)** Schematic of DNA curtains assay to test interaction between Reb1 and INO80 on DNA. **(F)** Same as (B), but for INO80 (cyan) co-localizing with Reb1 (yellow). **(G)** Top: Schematic showing possible outcomes of co-localization events. Bottom: Pie charts indicate fraction of co-localization events that result in a block for nucleosomes and Reb1.

After injection, we observed INO80 co-localizing with nucleosomes over extended periods with a median interaction time of 15.3 *±* 4.6 s (fig. 5B,C; fig. S5A). Closer analysis of co-localization times revealed three types of interactions (fig. S5B). First, extremely short co-localizations (0.23 *±* 0.10 s, 24.5 %), which most likely constitute collisions without nucleosome binding. Second, short associations (16.6 *±* 3.2 s, 41.3 %) that were represented by complexes binding dynamically to nucleosomes in-between 1D searches. These interaction times are in line with previously reported nucleosome occupancy times of other remodelers [27]. Third, we also detected a cohort of extremely long-lived co-localizations (1200 *±* 660 s, 34.1 %) that were mostly characterized by complexes already bound to a nucleosome at the start of recording and that remained statically bound until they dissociated or photobleached (fig. S5A). 85.3 % of INO80 molecules (29/34), for which we could determine the method of association, associated with the nucleosome through 1D searches, as opposed to 3D searches (fig. S5C).

Contrasting a previous study of RSC and ISWI [27], INO80 did not induce long-range nucleosome translocation, most likely because remodeling distances were below our resolution limit. To ensure that the used INO80 complex was active even in its labeled form, we confirmed the remodeling capability with gel-based sliding assays (fig. S5D).

### No indication of H2A.Z/H2B-dimer eviction by INO80

Specific eviction of H2A.Z-histones by INO80 *in vivo* and *in vitro* has been a controversial topic in INO80 research, with some studies detecting radical changes in H2A.Z distribution across the genome upon INO80 deletion [47], while others observe no differences [48]. Labeling of the H2B-histone allowed us to observe whether INO80 evicts H2A.Z/H2B-dimers from *λ*-DNA during our assay. While we could not distinguish whether steps in fluorescent signals were caused by photobleaching or H2A.Z/H2B eviction, we were able to compare the time until fluorophore signal disappearance of nucleosomes bound by INO80, with those of nucleosomes imaged in absence of INO80. Both were well-described by a single exponential distribution with a mean lifetime of 193.0 *±* 23.1 s and 244.0 *±* 28.6 s, respectively (fig. 5D). The disappearance rate between both categories showed no significant difference (*p* = 0.29, log-rank test), indicating that INO80 did not substantially enhance the eviction rate for H2A.Z/H2B-dimers in our assay.

### INO80 does not stably bind to DNA-bound Reb1

Apart from DNA sequence, nucleosome positioning by INO80 is influenced by general regulatory factors (GRFs), which bind DNA at defined motif sites and have been proposed to act as specific barriers to the remodeling process and thereby shape nucleosome positioning [7, 20]. Whether this type of regulation requires INO80 binding to GRFs or whether these proteins just act as passive roadblocks for remodeling is unclear. To answer these questions we purified and labeled the GRF Reb1 from *S. cerevisiae* and assessed its integrity on DNA curtains. Purified Reb1 was functional and bound to known stong binding sites on *λ*-DNA (fig. S5E). To test whether INO80 and Reb1 interact on DNA, we loaded both proteins in consecutive steps onto DNA curtains (fig. 5E). We did not observe binding of INO80 from solution to DNA-bound Reb1. Additionally, we did not observe any simultaneous binding of both proteins. This was unchanged when we reversed the order of injections or co-incubated INO80 and Reb1 for five minutes prior to injection. We then analyzed interactions between INO80 and Reb1 arising through encounters during INO80 1D search (fig. 5F). Co-localization times were extremely short with only 13 % of associations being longer than 2 s and a median of 0.80 *±* 0.19 s, suggesting that INO80 does not specifically interact with DNA-bound Reb1 for extended periods of time (fig. 5C). Hence, we conclude that Reb1 functions as a passive barrier for INO80 during nucleosome positioning.

### Nucleosomes and DNA-bound Reb1 act as barriers for INO80 diffusion

Finally, to understand how INO80 locates the promoter and the +1 nucleosome within a crowded chromatin environment, it is crucial to uncover how INO80’s path along DNA during 1D searches is affected by collision or interaction with other proteins. We therefore asked whether nucleosomes or Reb1 might act as roadblocks for INO80, confining the complex within subsegments of the promoter. Accordingly, we divided all interactions of INO80 with nucleosomes and Reb1 that showed 1D diffusion before and after, into *block* and *bypass* events (fig. 5G). *Blocks* were defined as events where INO80 stays on the same side of the interaction partner; all remaining interactions were defined as *bypasses*. INO80 was clearly confined by both proteins with 90 % of nucleosome (81*/*90) and 98 % of Reb1 interactions (51*/*52) resulting in a *block*. We then explicitly looked at longer co-localizations with nucleosomes (*>*5 s), as in these cases INO80 interacts unequivocally with nucleosomes and passive collisions are excluded (fig. S5F). These events showed a similar picture with 86 % (25*/*30) of interactions categorized as *blocks*. This indicates that nucleosomebound INO80 preferably disengages to the same direction had bound from and does not hop over the nucleosome. Together, these results suggest that INO80 diffusion is blocked by both collisions and interactions with nucleosomes and Reb1. Additionally, these observations validate our earlier findings that INO80 mostly uses a sliding mechanism to diffuse along DNA.

## DISCUSSION

### DNA binding and promoter targeting

Depending on the measurement method each *S. cerevisiae* cell nucleus harbors 57 000 to 60 000 nucleosomes [49]. Of these, only 4500 to 5500 (7.5 to 9.6 %) constitute possible +1 nucleosomes [48, 50], the main target of INO80 [14]. A unique trait highlighting +1 nucleosomes is the unilateral presence of long stretches of free neighboring promoter DNA. Our experiments clearly demonstrate that INO80 binds strongly to DNA and prefers furthermore certain DNA shape properties, e.g. high propeller twist (Ψ) and high electrostatic potential (EP) (fig. S2G). Here, INO80 exhibits a notably increased on- and reduced off-rate, resulting in accumulation of the complex on these sequences (fig. 2G). Intriguingly, TATA-less promoter and ORI DNA in *S. cerevisiae* is greatly enriched in poly(dA:dT) sequence tracts that are characterized by especially high Ψ and high EP [28, 51]. Closer inspection poly(dA:dT)-tracts in promoter sequences revealed further that they are positioned next to each bordering nucleosome and that their strength positively correlates with direction of transcription [10, 28] (fig. 6A). Given the strong shape preferences we observed by INO80, these asymmetrically positioned poly(dA:dT) tracts could guide the complex towards promoters in general and towards +1 nucleosomes in particular and explain how differential recognition of +1 and *−*1 nucleosomes is achieved. In general, INO80 shows a high preference for long linkers in nucleosome binding and remodeling assays [18, 40, 41, 45], which can be explained by its structure. With the N- and A-module the complex possesses two DNA-binding modules, which bind extranucleosomal DNA in tandem with a combined footprint of *≈*70 bp [14]. Linkers within gene bodies, on the other hand, are short in *S. cerevisiae* with a length of only *≈*18 bp [52].

**FIG. 6.**
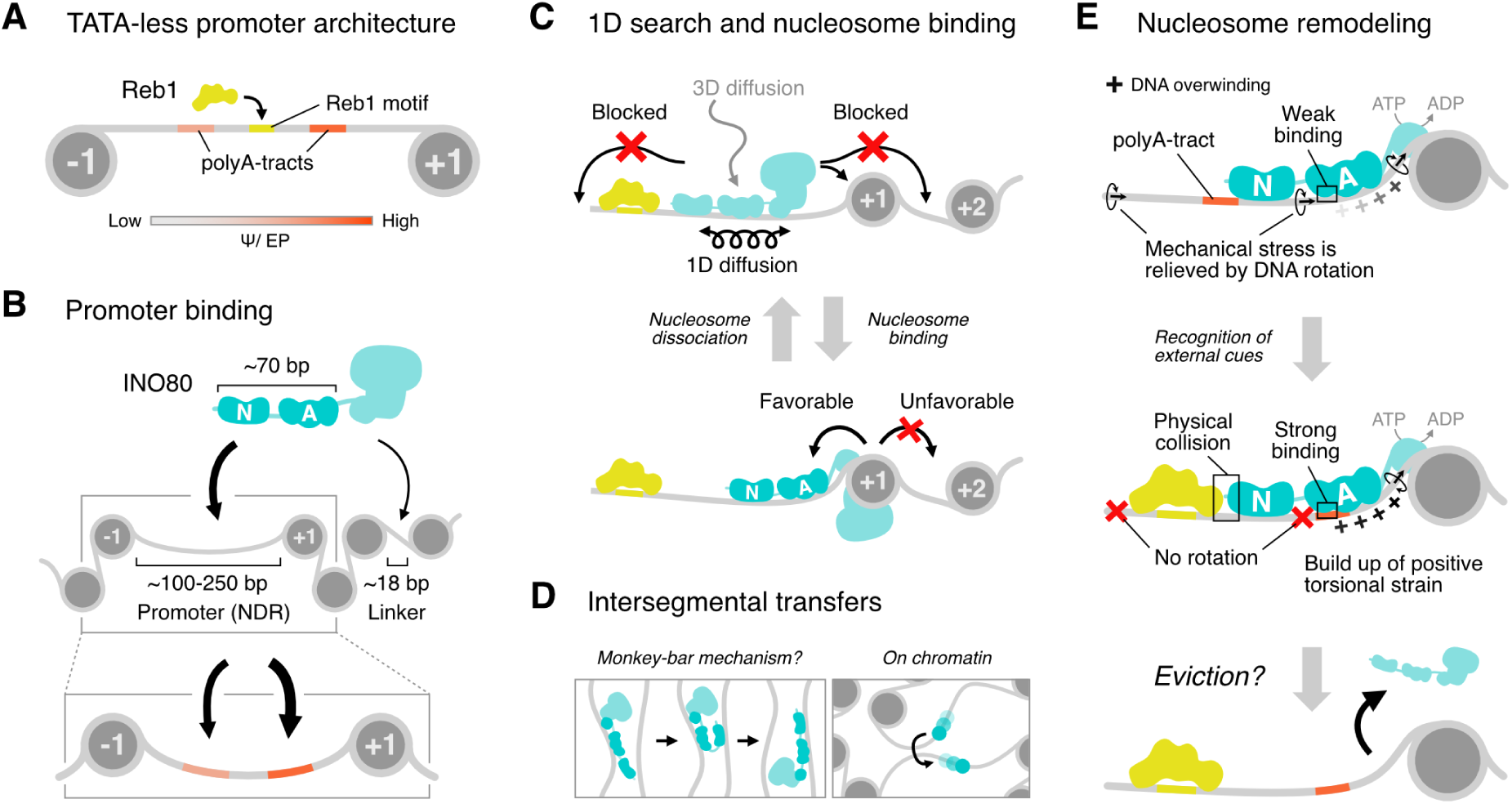
Model for INO80 target site-search and DNA-shape dependent remodeling regulation. **(A)** Scheme of promoter architecture of TATA-less genes in *S. cerevisiae*. Reb1 motifs (yellow) are found in the center of the NDR, while poly(dA:dT)-tracts (orange) are positioned *≈*40 to 55 bp away from the bordering nucleosomes. Poly(dA:dT)-tracts are characterized by high Ψ and EP and show stronger enrichment next to +1 nucleosomes. **(B)** INO80 is recruited to promoters through its strong preference for long free DNA stretches and read-out of DNA shape properties. Linker DNA in gene bodies on average is too short for proper binding of INO80s DNA-binding modules (dark teal). The positioning of preferred DNA shapes guides INO80 into the vicinity of the +1 nucleosome. **(C)** Once bound to the promoter, INO80 performs 1D searches to associate with the +1 nucleosome. Due to its sliding mechanism, Reb1 (yellow) and nucleosomes act as roadblocks to INO80 diffusion and confine it within the promoter. Placement of its DNA-binding modules strongly disfavors dissociation from the nucleosome into the gene body. **(D)** During 1D searches INO80 can transfer between DNA strands. This might be possible due to its multiple DNA-binding domains which are flexibly connected and would allow bridging of DNA strands in dense chromatin through a “monkey-bar mechanism”. **(E)** During remodeling, DNA is translocated in a corkscrew-motion towards the nucleosome by the Snf2-type motor, generating overwound DNA in front of the motor, which is relieved by free rotation of the DNA. Simultaneously the extranucleosomal DNA is probed by the upstream A- and N-module. Strong binding of the A-module to an A-tract or physical collision of the N-module with Reb1 lead to fixation of the DNA. As the motor continues to translocate DNA, positive torsional strain is build up, which ultimately might cause eviction of the complex.

Combined, these findings are consistent with a model where steric restrictions prevent INO80 from proper binding within the gene body (fig. 6B). In consequence, INO80 is targeted to NDRs through recognition of poly(dA:dT)-tract signals within yeast promoters and ORIs and thereby guided into proximity of flanking nucleosomes. The fact that poly(dA:dT)-tract enrichment is mostly missing in promoters with a TATA-box, might imply a heightened importance for +1nucleosome positioning by INO80 at TATA-less promoters. Indeed, TATA-less promoters show more uniform +1 nucleosome positioning and recent studies indicate that in absence of a canonical TATA-box, effective assembly of the pre initiation complex is reliant on the position of the +1 nucleosome [53].

### 1D search at a crowded promoter

Once INO80 is bound at an NDR, it needs to associate with the neighboring +1 nucleosome. Here, we observed that INO80 performs 1D diffusion along DNA (fig. 1A–D, fig. 2A) and presented examples of dynamic interactions with H2A.Z-containing nucleosomes (fig. 5B). It is well-established that nuclear proteins, which act on DNA or DNA-bound targets, combine diffusion in 3D space and 1D along DNA to reduce search times for their targets in a process called ‘facilitated diffusion’ [23, 33, 54–56]. The majority of interactions we recorded were initiated directly through 1D searches (fig. S5C), showing that INO80 interactions with +1 nucleosomes through prior 1D searches represent a possible pathway of association *in vivo*. Our observations therefore reinforce the notion that chromatin remodeling complexes use facilitated diffusion, which is supported by other recent single-molecule studies with chromatin remodelers RSC, IWS2 and SWR1 [26, 27]. Note that we cannot exclude that the ratio of 1D to 3D association might be biased towards 1D association, due to the scarcity of nucleosomes on our DNA templates.

By closer analysis of 1D diffusion kinetics, we found that INO80 diffuses mostly by sliding along the helical pitch of the DNA (fig. 2A,D). In line with this, nucleosomes and the GRF Reb1 acted as roadblocks for INO80 movement (fig. 5B,F,G). Additionally, we observed that INO80 continued 1D searches after interacting with nucleosomes and preferably dissociated back into the same direction it bound from (fig. S5F). This preference can be explained by the tandem placement of its DNA-binding modules on the entry site of the extranucleosomal DNA. Dissociation into the opposite direction would require big conformational changes, i.e. un- and rebinding of both modules onto the DNA on the exit side of the nucleosome. Not being able to bypass nucleosomes, even by dissociation after remodeling, could capture INO80 within the NDR and prevent “spilling” into intragenic regions (fig. 6C). Coupled with poly(dA:dT) tract-mediated DNA binding, this mechanism might drive differential +1 nucleosome remodeling by INO80. In addition, it promotes repositioning of nucleosomes into the NDR, by assuring that INO80 binds them in the correct orientation.

### Intersegmental transfers

In areas with high DNA density, we repeatedly observed INO80 and *Ct.* ΔN-INO80 complexes translocating to a neighboring DNA strand without changes in *y*-position (fig. S2A–D, fig. S3D), indicating the occurrence of intersegmental transfers. Intersegmental transfers have been shown to increase efficient target site search in theoretical models and in single-molecule experiments for small DBPs by reducing local oversampling, especially in environments with high DNA density, where many transfer opportunities arise [24, 57, 58]. It is tempting to speculate that to achieve this, INO80 might—like ISWI—use a “monkey-bar” mechanism to transfer to the neighboring DNA-strand, a bridging mechanism that would be available to protein complexes with multiple DNA-binding domains/subunits that are flexibly connected [59, 60] (fig. 6D). In addition, all INO80 variants were able to jump in *cis*, most likely through dissociation and reassociation onto the same DNA (fig. S2E, fig. S3G). Enhancement in mobility through intersegmental transfers could increase overall +1 nucleosome targeting, especially in euchromatic regions or in the context of transcriptional condensates, where NDRs are thought to be in close proximity [61].

### Implications for nucleosome remodeling

After association with the +1 nucleosome, INO80 pulls it towards the NDR [5]. For accurate positioning, INO80 remodeling has to be stopped through negative regulation by external cues. So far, two such cues have been identified indirectly, through analysis of nucleosome positions *in vitro* and *in vivo*. The first cue includes blocking of DNA translocation by a barrier protein, bound to DNA (e.g. Reb1), through an unknown mechanism [7, 10], while the second involves DNA shape features within the nucleosomal and extranucleosomal DNA, which impede the remodeling process [10, 21]. Note, that promoters with Reb1 motif also contain positioned DNA shape features and current data indicates a synergizing effect of both cues.

Inhibition through a physical barrier might include specific interactions between the barrier and INO80 through protein-protein-contacts with the N-module, which inhibit INO80’s remodeling capabilities allosterically. Alternatively, a passive barrier bound strongly to DNA could be enough to prevent INO80 from reeling in further DNA and specific contacts would not be required. We showed that Reb1, bound to its motif, is a potent barrier for INO80 diffusion, but strikingly, we did not observe long-lived interactions that imply specific binding (fig. 5E–G). Therefore, our observations are consistent with the passive barrier model. In line with this, Reb1 can act as a regulator for both +1 nucleosome positions at bidirectional promoters *in vivo*, and both ORC and Reb1 can perform this function irrespective of motif orientation [7, 10]. Furthermore, to our knowledge, no common interaction motifs between INO80 and any of the proposed barrier proteins have been found so far.

Inhibition through DNA shape features on the other hand requires tracking and read-out of DNA shape by INO80 during nucleosome remodeling. *Ct.* ΔN-INO80, which represents the minimally required complex for functional and competent nucleosome positioning, displayed potent DNA shape-readout during DNA scanning (fig. 3C,E, fig. S3B,C), demonstrating that DNA shape modulates INO80 binding properties dynamically during the scanning process. Further, we observed that the A-module can individually recognize high Ψ DNA and is most likely responsible for recognition of structural DNA features by INO80 (fig. 3E, fig. S3E,F). In agreement with this, we showed that recognition of a high-Ψ motif within linker DNA by the A-module can lead to abortion of the remodeling process (fig. 4A–D). This observation is consistent with DNA shape profiles of +1 nucleosomes, remodeled by INO80 *in vitro*, which show high Ψ values at *−*110 to *−*125 bp from the nucleosome dyad, where the N-terminal end of the A-module is expected to bind the DNA [10]. Furthermore, poly(dA:dT)-tracts at TATA-less genes *in vivo* are positioned at the same distance relative to the +1 nucleosome dyad [28]. While the molecular details of DNA shape recognition remain unclear, the set of shape features preferred by INO80 imply a readout of the negative electrostatic potential within the minor groove through arginines and lysines (fig. S2G) [37]. This is supported by the findings of a recent study, which showed that mutation of four positively charged amino acids at the N-terminal end of the A-module lead to a severe speed-up of remodeling and ATP hydrolysis, while drastically reducing the DNA binding capabilities of the A-module [19].

Taken together, these findings suggests that inhibition of remodeling by INO80 through DNA shape is connected to stronger binding by the A-module. Interestingly, this indicates that remodeling inhibition through a barrier or through DNA shape features may act by a similar mechanism, namely by fixing the DNA at a certain position in front of the ATPase domain. Although anchoring of the DNA provides a mechanism to prevent effective DNA translocation, it seems counterproductive for dissociation of the complex. Intriguingly, current models predict overwinding of the DNA in front of the motor during remodeling [42]. Strong binding of the A-Module might enhance this effect: as the motor goes through additional ATP-cycles, the deformation stress of the DNA cannot be resolved by rotation anymore (fig. 6E). This would cause additional positive torsional strain, which is associated with minor groove narrowing [62] and could therefore stabilize A-module-DNA interactions even further. However, whether these twist deformations could directly cause eviction of the complex by disrupting the binding of the ATPase motor or whether they might induce conformational changes that lead to eviction has to be investigated by future research.

### Conclusion

Our results demonstrate that both the target-site search of INO80 and its capacity to accurately position nucleosomes are closely connected to its ability to dynamically read out DNA shape features. Intriguingly, the remodeling activities of others remodelers like ISW1a or RSC are also modulated by these features [63]. In light of these findings, DNA shape patterns, originating through poly(dA:dT)-tracts, emerge as central cues, around which remodeler recruitment, activity and therefore promoter architecture is organized, opposing previous theories of intrinsic nucleosome exclusion mechanisms. Additionally, our observations underpin recent studies, which show that the concept of facilitated diffusion extends to chromatin remodelers. In the case of INO80, 1D and 3D diffusion are combined with characteristics of its molecular architecture to exploit traits that highlight target nucleosomes and thereby maximize engagement. Taken together, our study supports the notion that remodelers have evolved into complex machines that are able to simultaneously integrate multiple forms of information to construct diverse chromatin landscapes.

## MATERIALS AND METHODS

### Protein expression and purification

#### S.cerevisiae Wild-type INO80

The yeast strain for *S. cerevisiae* Wild-type INO80 originated from the S288c genetic background and was generated as previously described [64]. A 3x-FLAG-tag and a ybbR-tag connected by a short linker were chromosomally inserted at the C-terminus of Ino80 using pBP83 as a template.

Wild-type INO80 was expressed endogenously from *S. cerevisiae* and purified similar to a previously described protocol [64]. W303-1a cells were grown in 8 L YPD for 24 h at 30 ^◦^C and were then collected by centrifugation (6000 g, 10 min, 4 ^◦^C). The pellet was resuspended in an equal volume of 2x lysis buffer (1x = 25 mM HEPES-KOH pH 7.6, 500 mM KCl, 10 % glycerol, 0.05 % NP-40, 1 mM EDTA, 1 mM DTT, 4 mM MgCl_2_) with protease inhibitors 0.2 mM PMSF (Roth), 0.2 µM pepstatin A (Genaxxon), 0.2 µg*/*mL aprotinin (Genaxxon), and 2 µM leupeptin (Genaxxon). Cells were then frozen in liquid nitrogen in a dropwise manner and the frozen cells were crushed using a Freezer Mill (SPEX SamplePrep 6875 Freezer/Mill) (6 cycles for 2 min, crushing rate 15). For purification of the complex, the frozen cell powder was thawed, resuspended in 50 mL 1x lysis buffer with protease inhibitors and cleared by ultracentrifugation (235 000 g, 1 h, 4 ^◦^C). The supernatant was incubated with 1.5 mL pre-washed anti-FLAG M2 affinity gel resin (Sigma) for 1 h at 4 ^◦^C and applied to a gravity flow column. The resin was washed with 4 column volumes (CV) of lysis buffer and 1 CV of wash buffer 25 mM Tris-HCl pH 7.2, 200 mM KCl, 10 % glycerol, 0.05 % NP-40, 1 mM EDTA, 1 mM DTT, 4 mM MgCl_2_). INO80 was eluted by 1 mL of the same buffer with 0.5 mg*/*mL 3xFLAG peptide (Sigma), followed by 2x 1 mL of buffer with 0.25 mg*/*mL 3xFLAG peptide. Eluted fractions were analyzed by SDS-PAGE, and pooled.

For fluorescent labeling, Sfp (made in-house) was mixed with INO80 in a molar ratio of 2:1 and a 10x excess of LD555-CoA dye (Lumidyne) as well as 15 mM MgCl_2_. The reaction was incubated for 30 min at 25 ^◦^C and further purified with a Mono Q 5/50 GL column using a gradient from 100 mM to 600 mM KCl (25 mM Tris-HCl pH 7.2, 25 % glycerol, 0.05 % NP-40, 1 mM EDTA, 1 mM DTT, 4 mM MgCl_2_). The protein was stored at *−*70 ^◦^C.

The unlabeled *S.cerevisiae* wild-type INO80 complex used for sliding assays was purified as previously described [10].

#### C. thermophilum Ct. ΔN-INO80

The *Ct.* ΔN-INO80 mutant from *C. thermophilum* was cloned using the MultiBac technology. The gene coding for Ino80^718–1848^–2xFLAG-ybbR was cloned in pACEBac1; genes for Rvb1 and Rvb2 were cloned in pIDC; and genes coding for Arp5, Ies6, and Ies2 were cloned in the pIDK vector. Together, they were combined in one bacmid. Genes coding for Ies4 and Taf14 were cloned in pACEBac1, and genes coding for Arp8, actin, and Arp4 were cloned in a pIDK vector and combined on a separate bacmid. PirHC (Geneva Biotech) and *E. coli* XL1-Blue (Stratagene) cells were used for all recombination steps by the addition of Cre recombinase (NEB). Both baculoviruses (1:200) were used to infect 1 L of Trichoplusia ni High Five cells (Invitrogen, B85502) for expression.

After addition of viruses, cells were cultured for 60 h at 27 ^◦^C and harvested by centrifugation at 4 ^◦^C. Cells were disrupted in lysis buffer (30 mM HEPES pH 7.8, 400 mM NaCl, 10 % glycerol, 0.25 mM dithiothreitol (DTT), 0.28 µg*/*mL pepstatin A, 0.17 mg*/*mL phenylmethylsulfonyl fluoride (PMSF), 0.33 mg*/*mL benzamidine, 2 mM MgCl_2_) for complex purification and gently sonicated. Raw lysate was cleared by centrifugation at 20 500 g at 4 ^◦^C for 30 min. Supernatant was incubated with 2 mL of ANTI-FLAG M2 Affinity Gel (Sigma-Aldrich) for 1 h and washed with 50 mL of lysis buffer and 75 mL of wash buffer (30 mM HEPES pH 7.8, 150 mM NaCl, 5 % glycerol, 1 mM DTT). The protein was eluted by incubation with 4.5 mL of wash buffer (supplemented with 0.2 mg*/*mL FLAG peptide) in three incubation steps of 20 min each and the elution fractions were concentrated.

For ybbR-tag labeling, Sfp (made in house) was mixed with *Ct.* ΔN-INO80 in a molar ratio of 4:1 and a 5x excess of LD555-CoA dye (Lumidyne) as well as 10 mM MgCl_2_. Reaction was performed for 40 min at 25 ^◦^C. The labeling reaction was loaded onto a Mono Q 5/50 GL column (GE Healthcare) and eluted by an increasing salt gradient (0.2 to 1 M NaCl), resulting in highly pure labeled *Ct.* ΔN-INO80.

#### S. cerevisiae INO80 A-module

The *S.cerevisiae* INO80 A-module mutant was cloned using the MultiBac technology. The gene coding for Ino80^440-598^–2xFLAG-ybbR was cloned into pLIB by Gibson assembly of PCR products containing the ybbR-pLIB backbone and Ino80^440-598^–2xFLAG. The coding sequences of the INO80 subunits comprising the A-module (Arp4, Arp8, N-Actin, Ies2, Ies4, Taf14) were cloned into pFBDM vectors and then combined into a single vector. All cloning was carried out in *E. coli* XL1-Blue (Stratagene). From each bacmid (generated in *E. coli* DH10 MultiBac cells), baculoviruses were generated in Spodoptera frugiperda (Sf21) insect cells (Thermo Fisher Scientific, 11497013). Both baculoviruses (1:200) were used to infect 100 mL Trichoplusia ni High Five cells (Invitrogen, B85502) for expression.

After addition of virus, cells were grown for 70 h and harvested by centrifugation at 4 ^◦^C. All further steps were performed at 4 ^◦^C. After resuspension in lysis buffer (50 mM Tris-HCl pH 7.9, 500 mM NaCl, 10 % glycerol, 2 mM DTT), cells were supplemented with half a protease inhibitor tablet, lysed by sonication (duty cycle 40 %, Intensity 5) and centrifuged for 30 min at 30 000 g. Lysate was incubated with 100 µL anti-FLAG M2 affinity gel resin (Merck, A2220) for 1 h while rotating, before supernatant was removed by centrifugation (5 min, 1000 g). Anti-FLAG beads were then washed twice with 1 mL Wash buffer A (25 mM HEPES pH 8.0, 500 mM KCl, 10 % glycerol, 0.05 % IGEPAL CA630, 1 mM DTT, 4 mM MgCl_2_) and twice with Wash buffer B (25 mM HEPES pH 8.0, 200 mM KCl, 10 % glycerol, 0.05 % IGEPAL CA630, 1 mM DTT, 4 mM MgCl_2_). The protein was eluted in two steps by addition of 200 µL Wash buffer B (supplemented with 0.2 mg*/*mL FLAG peptide) for 20 min.

For labeling, the eluate was supplemented with MgCl_2_ to a final concentration of 15 mM and mixed with Sfp (made in-house) in a molar ratio of 2:1 and a 10x excess of LD555-CoA dye (Lumidyne). The reaction was incubated for 30 min at 25 ^◦^C and further purified by size exclusion chromatography using a Superdex S200 10/300 GL (Cytiva) in Wash buffer B. The protein was stored at *−*70 ^◦^C.

#### S. cerevisiae Reb1

The pET21-Reb1 construct with a N-terminal 10x-Histidine-tag and a ybbR-tag connected by a short linker was generated by Gibson assembly of three PCR-generated fragments containing the pET21 backbone, both tags and the gene respectively. All cloning for the construct was carried out in *E. coli* XL1-Blue (Agilent Technologies).

For expression, 500 mL culture of *E. coli* Rosetta(DE3) (Merck) was grown in LB medium, induced with 200 µM IPTG, cultivated for 4 h at 37 ^◦^C and harvested by centrifugation at 4 ^◦^C. All further purification steps were performed at 4 ^◦^C. Cells were then resuspended in lysis buffer (50 mM NaH_2_PO_4_ pH 8.0, 100 mM NaCl, 20 mM imidazole, 10 % glycerol, 2 mM MgCl_2_) supplemented with a protease inhibitor tablet. Cells were then lysed by sonication (duty cycle 40 %, Intensity 5) and 0.1 µL*/*mL Pierce universal nuclease was added. After 30 min the lysate was cleared by centrifugation (20 min, 20 000 g) then 5 mM beta-mercaptoethanol was added and salt concentration was adjusted to 500 mM NaCl. A gravity flow column was prepared with 2 mL of Ni-NTA beads (Macherey-Nagel) equilibrated with 10 CV of lysis buffer and the supernatant was added in three steps of 20 min. Afterwards the beads were washed twice with 10 CV of wash buffer (50 mM NaH_2_PO_4_ pH 8.0, 1.5 M NaCl, 40 mM imidazole, 10 % glycerol, 2 mM MgCl_2_, 5 mM beta-mercaptoethanol). The protein was eluted from the column with 7.5 CV of elution buffer (50 mM NaH_2_PO_4_ pH 8.0, 150 mM NaCl, 250 mM imidazole, 10 % glycerol, 2 mM MgCl_2_, 5 mM beta-mercaptoethanol). Fractions containing Reb1 were pooled and dialyzed against Heparin buffer (25 mM HEPES pH 7.5, 150 mM NaCl, 5 % glycerol, 1 mM DTT) for 16 h before further purification on a Cytiva HiTrap Heparin HP column with a 150 to 1000 mM NaCl gradient.

For labeling, the eluate was supplemented with MgCl_2_ to a final concentration of 15 mM and mixed with Sfp (made in-house) in a molar ratio of 2:1 and a 10x excess of LD655-CoA dye (Lumidyne). The reaction was incubated for 30 min at 25 ^◦^C and further purified by size exclusion chromatography using a Superdex S200 10/300 GL (Cytiva) in Reb1 storage buffer (25 mM HEPES pH 7.5, 200 mM NaCl, 10 % glycerol, 1 mM DTT). The protein was stored at *−*70 ^◦^C.

#### SpyCatcher003

A pET28a-based plasmid containing a SpyCatcher003 construct with N-terminal MGGGC- extension and C-terminal 6xHis-tag was transformed in Rosetta (DE3). Cells were grown at 37 ^◦^C to optical density 600. The expression was induced by 0.4 mM IPTG and carried out at 18 ^◦^C overnight. Harvested cells were resuspended in lysis buffer (25 mM Tris-HCl pH 7.5, 500 mM NaCl, 15 mM imidazole) supplemented with 1 mM PMSF and 0.1 mg*/*mL lysozyme, sonicated, and centrifuged. The supernatant was loaded onto a HisTrap FF-columns (GE Healthcare, Chicago, IL), washed with 10 CV lysis buffer, and eluted in lysis buffer containing 200 mM imidazole. Pooled fractions were gel filtrated using a Sephacryl S300 (GE Healthcare) or Superdex S200 (GE Healthcare) column. Eluted proteins were equilibrated in storage buffer (25 mM Tris-HCl pH 7.2, 150 mM NaCl) and stored at *−*70 ^◦^C.

#### S. pyogenes dCas9

To create the expression construct, the coding sequence for nuclease deficient dCas9 was combined with an N-terminal Spytag002 and a C-terminal intein-CBD (chitin-binding-domain) cassette in a pet28a-vector by Gibson assembly. All cloning for the construct was carried out in *E. coli* XL1-Blue.

*S.pyogenes* dCas9 was expressed in *E. coli* Rosetta(DE3) (Merck). 2 L culture was grown in LB medium, induced with 200 µM IPTG, cultivated for 24 h at 18 ^◦^C and harvested at 4 ^◦^C. Pellets were resuspended in 50 mL loading buffer (25 mM Tris pH 7.5, 150 mM NaCl) supplemented with 1 mM PMSF and 20 µg*/*mL lysozyme and cells were then lysed by sonication (duty cycle 40 %, Intensity 5). The lysate was cleared by centrifugation (30 min, 2820 g, 4 ^◦^C) and added to 4 mL equilibrated chitin resin (NEB IMPACT-system) in a gravity flow column. The column was washed with 20 CV of wash buffer (25 mM Tris pH 7.5, 1 M NaCl) and on-column cleavage was induced by incubation in 3 CV wash buffer supplemented with 50 mM DTT for 40 h. To elute the protein, flow was allowed to continue, eluted fractions containing the protein were dialyzed against storage buffer (25 mM Tris pH 7.5, 150 mM NaCl, 1 mM DTT) and concentrated using Amicon Ultra-15 filters (30 kDa MWCO, Merck). The protein was stored at *−*70 ^◦^C.

The protein was labeled using the Spytag/Spycatcher system [65]. Spycatcher003 protein with an N-terminal MGGGC-extension was labeled by slow addition of a 4x molar excess of ATTO643-maleimide dye (ATTO-Tec) and rotation for 2 h at 25 ^◦^C. Precipitates were removed by centrifugation and excess dye by desalting using a HiTrap Desalting column (Cytiva) in Desalting buffer (100 mM Na_2_HPO_4_ pH 7.5, 150 mM NaCl). Labeled Spycatcher003 was then incubated with purified dCas9 for 30 min and the conjugation reaction was again cleaned up by size exclusion chromatography as described above, but in dCas9 storage buffer (10 mM Tris pH 7.5, 300 mM NaCl, 0.5 mM EDTA, 1 mM DTT, 10 % glycerol) and stored at *−*70 ^◦^C.

Purified dCas9 was incubated with annealed gRNA (Metabion) in ratio of 1:1.2 for 20 min at 25 ^◦^C and assembled RNP complexes were stored at *−*70 ^◦^C.

#### H. sapiens Histone octamer

Recombinant *H. sapiens* Histones H2A.Z and H4 were purchased (The Histone Source). To enable site-specific cysteine labeling of histone H2B, two mutations were introduced into the constructs of H2B and H3.2 by PCR and ligation (H2B: T116C, H3: C111A). All cloning for the constructs was carried out in *E. coli* XL1-Blue (Agilent Technologies).

*H. sapiens* histones H2A, H2B and H3.2 were expressed in *E. coli* LOBSTR (Kerafast). For each histone 1 L culture was grown in LB medium, induced with 400 µM IPTG, cultivated for 4 h at 37 ^◦^C and harvested at 4 ^◦^C. Cells were then resuspended in 10 mL Tris-sucrose buffer (50 mM Tris-HCl pH 7.5, 10 % (w/v) sucrose) and flash frozen at *−*80 ^◦^C. Histones were then purified from inclusion bodies and reconstituted to nucleosomes as described before, but with small adjustments [66]. 0.05 % (w/v) Tween-20 was used instead of Triton X-100. For the nucleosome reconstitutions the histone mixture was dialyzed against refolding buffer with a total of five buffer changes and refolding buffer was supplemented with 0.2 mM TCEP instead of beta-mercaptoethanol from the first buffer change onward. After dialysis, aggregates were removed by centrifugation and the mix was concentrated to 1.5 mL using Amicon Ultra-15 filters (10 kDa MWCO, Merck). The labeling reaction was started by slow addition of 10x molar excess of ATTO643-maleimide dye (ATTO-Tec) to the mixture and rotation for 16 h at 4 ^◦^C. Labeled octamers were purified from labeled dimers by size exclusion chromatography as described in the original protocol [66]. Purified canonical octamers were used for generation of mononucleosomes and H2A.Z-containing octamers for generation of chromatinized *λ*-DNA constructs.

#### H. sapiens Mononucleosome

6-FAM-labeled DNA template for 0N80 nucleosomes was produced by PCR, using a plasmid carrying the Widom 601 sequence, as well as 6-FAM labeled short primers and primers with overhangs, which generate the different linker DNA sequences. The PCR products were purified by anion exchange chromatography (CaptoQ) with a salt gradient from 150 mM to 2 M NaCl in TE-buffer (25 mM Tris-HCl pH 7.5, 0.2 mM EDTA) over 15 CV, followed by size exclusion chromatography using a HiPrep 16/60 Sephacryl S-300 HR column. The DNA was eluted in TE-buffer supplemented with 150 mM NaCl and was concentrated by ethanol precipitation. Concentrated DNA was then combined with *H. sapiens* histone octamer at a 1.1-fold excess in reconstitution buffer (25 mM HEPES pH 7.5, 2 M NaCl, 10 % glycerol, 0.25 mM DTT) in 7 kDa MWCO Slide- A-Lyzer Dialysis Units (Thermo Fischer). Using a pump the salt concentration was decreased to 50 mM NaCl over 18 h at 4 ^◦^C. After dialysis, the nucleosomes were purified by anion exchange chromatography using a SourceQ 1-ml column. Nucleosome species were eluted by a salt gradient from 450 to 800 mM NaCl over 25 CV in reconstitution buffer. Fractions containing nucleosomes were pooled and dialyzed against reconstitution buffer containing 50 mM NaCl at 4 ^◦^C overnight. Finally, nucleosomes were concentrated to 1 mg*/*mL using Amicon Ultra-0.5 mL filters (30 kDa MWCO, Merck) and stored at *−*70 ^◦^C. The construct linkers contained the following sequences:

*Standard*

5’ 601-TGCATGTATTGAACAGCGACCTTGCCTACCACCGTGTACTCGTTGCTCGATCCGATCGTTCGTTGCGTAGCTGGCAGTCG 3’

*SHL -12/13*

5’ 601-TGCATGTATTGAACAGCGACCTTGCCTACCACCGTGTACTCGTTGCTCGATCCGTTTTTTTTTTGCGTAGCTGGCAGTCG 3’

### Generation of *λ*-DNA constructs for DNA curtains

*λ*-DNA constructs for DNA curtains were prepared as previously described [67]. For the assembly of nucleosomes, *λ*-DNA constructs were concentrated using Amicon Ultra-15 filters (30 kDa MWCO, Merck). 30 µL of a 30:1 reconstitution mix (10 mM Tris-HCl pH 7.5, 1 mM EDTA, 2 M NaCl, 0.25 mM DTT, 10 % glycerol, 0.2 mg*/*mL BSA, 27 nM octamers^H2A.Z^, 0.9 nM *λ*-DNA) was dialyzed against reconstitution buffer (25 mM HEPES pH 7.5, 2 M NaCl, 10 % glycerol, 0.25 mM DTT) using 7 kDa MWCO Slide-A-Lyzer Dialysis Units (Thermo Fischer). The NaCl concentration was decreased to 50 mM over 16 h at 4 ^◦^C using a pump and the mix was dialyzed for an additional 4 h against reconstitution buffer containing 20 mM NaCl. The resulting chromatinized *λ*-DNA was centrifuged to remove aggregates and stored at *−*20 ^◦^C in 50 % glycerol.

### DNA curtains

#### Flow cell preparation

Custom made flow cells were assembled from silica-fused slides (UQG Optics) grafted with chromium barriers produced via E-beam lithography and cover slips with a double-sided tape as described previously [29].

Flow cell loading and lipid master mix preparation was adapted from a previous protocol [29]. In brief, 100 mg DOPC (Avanti 850375P-200mg) dissolved in 1 mL chloroform, 1 mL DOPE-PEG (Avanti 880130C-25mg) and 50 µL DOPE-biotin (Avanti 870273C-25mg) were mixed and stored at *−*20 ^◦^C. 100 µL Master Mix was dried using N_2_ followed by applying a vacuum for 16 h and subsequently resolved in 2 mL lipid buffer (10 mM Tris pH 7.5, 200 mM NaCl). Lipids were sonicated five times for 1 min with 1 min pauses on ice (amplitude 20 %, duty cycle 20 %), filtered through 0.22 µm PVDF filters and stored at 4 ^◦^C for 3 to 4 weeks. Lipids were diluted 1:10 in lipid buffer containing 20 mM MgCl_2_ and flow cells were incubated four times for 5 min with 200 µL diluted lipids. After washout, flow cells were incubated with 2 µL of 1 mg*/*mL anti-digoxigenin (produced in house) in 500 µL lipid buffer for 15 min, followed by 5 µL streptavidin (Carl Roth) in 1 mL BSA buffer (20 mM HEPES pH 8.0, 1 mg*/*mL BSA, 4 mM Mg(OAc)_2_). After this, 20 pM naked or chromatinized *λ*-DNA in BSA buffer was added in four steps with 5 min incubation time.

#### Microscope setup

DNA curtain experiments were carried out on a prism-type TIRF microscope (Nikon Eclipse Ti2), equipped with three illumination lasers (488, 561 and 640 nm Coherent OBIS), an electron multiplying charged coupled camera (iXon Life, Andor) and a syringe-pump-driven microfluidics system supplying the sample chamber. All experiments were carried out using an OptoSplit II (Cairn Research) with a HC BS 640 beam splitter (AHF Analysetechnik) and additional filters (585/65 BP + 700/75 BP, Thorlabs). Videos were recorded in NIS Elements (Nikon) and analyzed in Igor Pro 8 (Wavemetrics) using custom written code (see below). Videos were aquired with a frame rate and illumination time of 50 ms for all experiments with the exception of *Ct.* ΔN-INO80, where 200 ms were used.

#### Single-protein diffusion

For single-protein diffusion assays on *λ*-DNA, the protein was diluted in 80 µL DNA curtains buffer and immediately injected into the flow cell with a constant flow of 0.3 mL*/*min. The flow was stopped when the protein concentration reached its peak and interactions were observed for up to 8 min per video. Protein was removed from *λ*-DNA with a 2 M NaCl wash before another injection. Single-molecule measurements were performed in DNA curtains buffer (20 mM HEPES pH 8.0, 1 mg*/*mL BSA, 4 mM Mg(OAc)_2_, 1 mM ATP, 1 mM DTT, 80 mM KGlu if not indicated otherwise). *Ct.* ΔN-INO80 was measured at 60 mM NaCl and data shown for wt-INO80 alone was measured at 120 mM KGlu. Protein concentrations injected varied between 125 to 250 pM for INO80 variants, 2 nM for dCas9 and 75 pM for Reb1.

#### Nucleosome bleaching, INO80–Reb1 and INO80–nucleosome interaction

To determine nucleosome bleaching rates, flowcells were prepared using chromatinized *λ*-DNA (see above). The flowcell was washed with 350 mM NaCl to remove partially formed nucleosomes or labeled histones bound to DNA and remaining nucleosomes were imaged until they had all bleached. For remodeler interaction experiments, INO80 was then flushed in and the flow was stopped before live-imaging was initiated.

For Reb1 interaction assays, INO80 and Reb1 were injected either sequentially or incubated for 10 min at a 10x concentration before dilution with DNA curtains buffer and injection into the flowcell. Concentrations used were the same as in single protein diffusion experiments.

### Mononucleosome sliding assays

To analyze INO80 sliding activity dependent on linker DNA shape, 50 nM *S.cerevisiae* INO80 was incubated with 100 nM of 6-FAM-labeled nucleosomes in sliding buffer (25 mM HEPES pH 8.0, 60 mM KCl, 7 % glycerol, 0.1 mg*/*mL BSA, 0.25 mM DTT) at 25 ^◦^C. The reaction was started by addition of 1 mM ATP complexed with 2 mM MgCl_2_ and stopped at different timepoints (1, 2, 5, 10, 20 and 60 min) by addition of Lambda DNA (0.15 mg*/*mL; NEB). An ATP-negative control was taken before the start of the reaction and treated like the last sample of the time series. Nucleosome species were separated by native polyacrylamide gel electrophoresis (PAGE) on a 3 to 12 % acrylamide bis-tris gel (Invitrogen) and visualized using a Typhoon imaging system (GE Healthcare).

Sliding assays to control for sliding activity of labeled endogenous *S.cerevisiae* INO80 were conducted with 2 nM INO80 and 100 nM of 6-FAM-labeled nucleosomes. The rest of the assay was performed as described above.

### Data analysis

#### Statistical analysis

Statistical analysis was performed using Igor Pro 8 (Wavemetrics). Respective type of test is stated in the main text, figure caption or methods section. Significance was set at *p <* 0.05 and indicated using the following nomenclature: *p >* 0.05 = ns.; *p <* 0.05= *; *p <* 0.01= **; *p <* 0.001= ***; *p <* 0.0001= ****. The results were reproducible and conducted with established internal controls. All samples that met proper experimental conditions were included in the analysis.

#### Single molecule tracking

Video files were analyzed using custom-written software in Igor Pro (Wavemetrics). Protein trajectories were localized using an algorithm inspired by DAOSTORM [68] and tracked using an in-house Markov chain Monte Carlo-based algorithm. Trajectories were manually screened for detections of particles that displayed exceedingly small (surface-bound) movement or unexpectedly bright or dim fluorescence (typically *<*10 %) and these trajectories were filtered out. Trajectories with less than 30 detections were also omitted from diffusion and binding analysis.

#### Diffusion analysis

Mean squared displacements (MSDs)

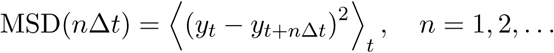

were determined from tracking data. The uncertainty of the MSD values were estimated using a published procedure [69] and used as weights for fits to the model

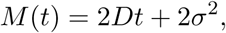

where *σ* is the localization noise. At short time delays, the MSDs are dominated by the relaxation time of the thermally fluctuating DNA, rather than movement of proteins along the DNA. Based on tracking of non-moving dCas9 molecules, the threshold where the DNA movement dominates was determined and MSD data with *t <* 200 ms were excluded from the fit. Fits were performed up to *n* = 100 *≡* 5 s for all proteins except *Ct.* ΔN-INO80, where *n* = 10 *≡* 2 s was used. Timevarying diffusion was identified by the same procedure applied to a sliding window of fixed length that was scanned along the trajectory. Errors for median diffusion coefficients are 68 % intervals from bootstrapping.

#### Calculation of the free-energy roughness

The Stokes-friction-dominated diffusion coefficient of a protein diffusing along DNA was modeled by

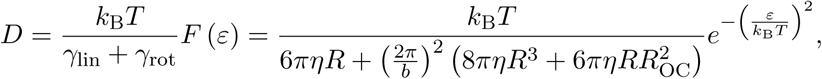

where *γ*_lin_ and *γ*_rot_ represent linear and rotational drag, *F* (*ε*) is the retarding factor due to an energy roughness *ε* (fig. 2B), *η* is the solvent viscosity, *b* is the distance between base pairs, *R* is the radius of the protein and *R*_OC_ is the distance of the protein from the DNA [32].

To determine *ε* for wildtype *S. cerevisiae* INO80, the median diffusion coefficient at 120 mM KGlu was used, *R* was estimated from cryo-EM structures to lie between 100 to 120 Å and *R*_OC_ was assumed to be the same as *R*.

To compare *ε* between wildtype *S. cerevisiae* INO80 and *Ch. thermophilum Ct.* ΔN-INO80, the median diffusion coefficient of both proteins at 40 mM KGlu and 60 mM NaCl respectively was used, *R* was estimated from cryo-EM structures to lie between 100 to 120 Å for wildtype *S. cerevisiae* INO80 and 95 to 115 Å for *Ch. thermophilum Ct.* ΔN-INO80 to account for the loss of the N-module. *R*_OC_ was assumed to be the same as *R*.

*ε* values of other DBPs were taken from previous single-molecule studies [32, 35].

#### DNA shape analysis and correlation with observables

The DNA propeller twist Ψ for each basepair was predicted using Deep DNAshape [38, 70] and averaged in 1 kbp bins.

The protein localization probability density along the DNA was calculated by summing all localizations within a bin and normalizing by the number of all localizations. Errors are 68 % intervals from bootstrapping. Binding events were binned according to the first detected localization of a trajectory, while mean lifetimes were binned according to the mean position of a trajectory along DNA. To determine the position-dependent apparent diffusion coefficient, *D*_App_ was calculated for all trajectories according to:

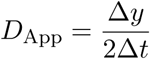

where the Δ*t* is a timeframe and Δ*y* is the measured displacement within that timeframe. For *Ct.* ΔN-INO80, Δ*t* was set to 2 s and for the A-module to 5 s. Calculated *D*_App_ values were binned depending on the mean position of *y* along the DNA. Pearson correlation coefficients were calculated by correlating the binned parameter with the binned mean Ψ values. Bins 0 to 5 kbp and 44 to 49 kbp were excluded from the correlations due to the proximity to the chrome barriers.

#### Interaction analysis

Interactions of INO80 with nucleosomes and Reb1 were determined manually from Kalmanfiltered tracks. Filtering was used to suppress the impact of fluctuations that are caused by movement of the DNA. To ensure that candidates for collisions are bound to the same DNA, the transversal movement perpendicular to the end-to-end vector of the DNA for both candidates was analyzed. Positive correlation and an cross-correlation decay compatible with the transversal relaxation time of the DNA was used as an indicator for binding to the same DNA. Interactions with 1D searches before and after the interaction were further separated into *block* if both 1D searches happened on the same side of the protein, or *bypass* otherwise. It should be noted that, using this definition, a bypass does not necessitate a jump of INO80 over the protein constituting a barrier, but can include specific binding to the protein followed by and 1D diffusion on the other side.

#### Eviction assay — Photobleaching

To assess H2A.Z/H2B dimer eviction by INO80, nucleosomes imaged with and without INO80 injection were tracked and the time durations until fluorescence signal loss were determined manually from fluorescence trajectories. Lifetimes of nucleosome fluorophores imaged without injection of INO80 were calculated and nucleosomes surviving beyond the end of the recording were marked as censored. For fluorophores of nucleosomes imaged with INO80, only the intervals during interaction with INO80 were considered and nucleosomes without bleaching/eviction events during the interaction were also marked as censored. Collected lifetimes were then compared using log-rank tests and survival plots were created using the Kaplan-Meier method [71]. Error bars are 68 % intervals from bootstrapping.

#### Sliding assay analysis

Band intensities were analyzed using ImageLab 6.1 (Bio-Rad) and IGOR Pro 8. Previous experiments have shown that hexasomes are remodeled by INO80 and that fully remodeled 0H80 hexasomes co-migrate with unremodeled 0N80 nucleosomes in native acrylamide bis-tris gels [12]. Nucleosome purification can contain trace amounts of hexasomes, visible as slight bands below the 0N80 nucleosome. To prevent erroneous calculation of the unremodeled nucleosome band, it was corrected for the amount of remodeled hexasome present at each timepoint. For that, the fraction of all nucleosome species including the hexasome were determined at every timepoint. Then, to determine how much hexasome was remodeled, the fraction of the hexasome band was compared to the first measured timepoint (1 min) and the difference was subtracted from the unremodeled nucleosome band (*U*). Finally, to determine whether nucleosomes with a poly(dA:dT)-tract at SHL *−*12/13 were remodeled more slowly, all nucleosome bands (unremodeled, intermediates, fully remodeled) were normalized to 100 % and fully remodeled (*R*) band fractions for different nucleosomes at each timepoint were compared using a two-tailed paired t-test. The paired replicates were performed in succession using the same solutions and vial of INO80 complex. The order of conditions was switched for each pair.

## ACKNOWLEDGMENTS

We thank Jonas Huber for initial experiments with nucleosomes and INO80 on DNA curtains. We thank Felix J. Metzner for sharing labeled *Ct.* ΔN-INO80, Franziska Därr, Annika Brehm, Mariia Likhodeeva and Alberto Ĺopez-Francos Ĺopez-Romero for guidance regarding nucleosome preparations and helpful discussions. We thank K.-P. Hopfner for providing INO80 plasmid constructs. We also thank Christoph Kurat, Priyanka Bansal and Silvia Haertel for help with expression and purification of endogenous *Sc.* INO80. Thanks to all members of the Stigler lab for helpful discussions and assistance throughout this work. J.S. acknowledges support by the LMU Center for Nanoscience CeNS.

## AUTHOR CONTRIBUTIONS

Investigation: GL, DH; methodology: GL; data analysis: GL, JS; data visualization: GL, SZ; programming: GL, JS; writing: GL, SZ, JS; supervision: JS; funding aquisition: JS.

## Supplementary Information

**FIG. S1.**
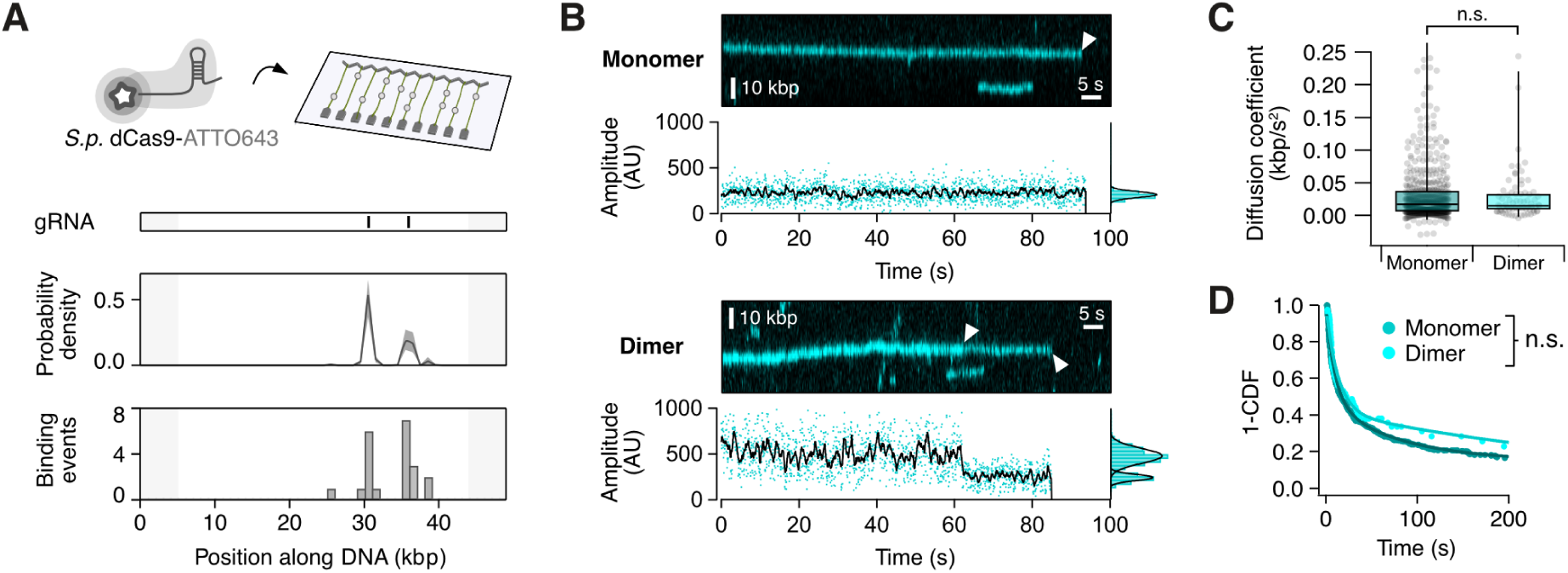
Photobleaching of monomeric and dimeric INO80 complexes. **(A)** Top: Schematic of dCas9 and DNA curtains assay. gRNA-binding motifs, probability density of localizations and binding events of dCas9 along *λ*-DNA. **(B)** Exemplary kymograms and fluorescence amplitudes of monomeric (top) and dimeric (bottom) INO80 complexes. **(C)** Comparison of diffusion coefficients of INO80 on *λ*-DNA for different multimeric states. Significance was determined using a Mann-Whitney U test. *n*_Monomer_ = 604. *n*_Dimer_ = 73. **(D)** Survival plot of INO80 on *λ*-DNA for different multimeric states. Rates were compared with Log-Rank test.

**FIG. S2.**
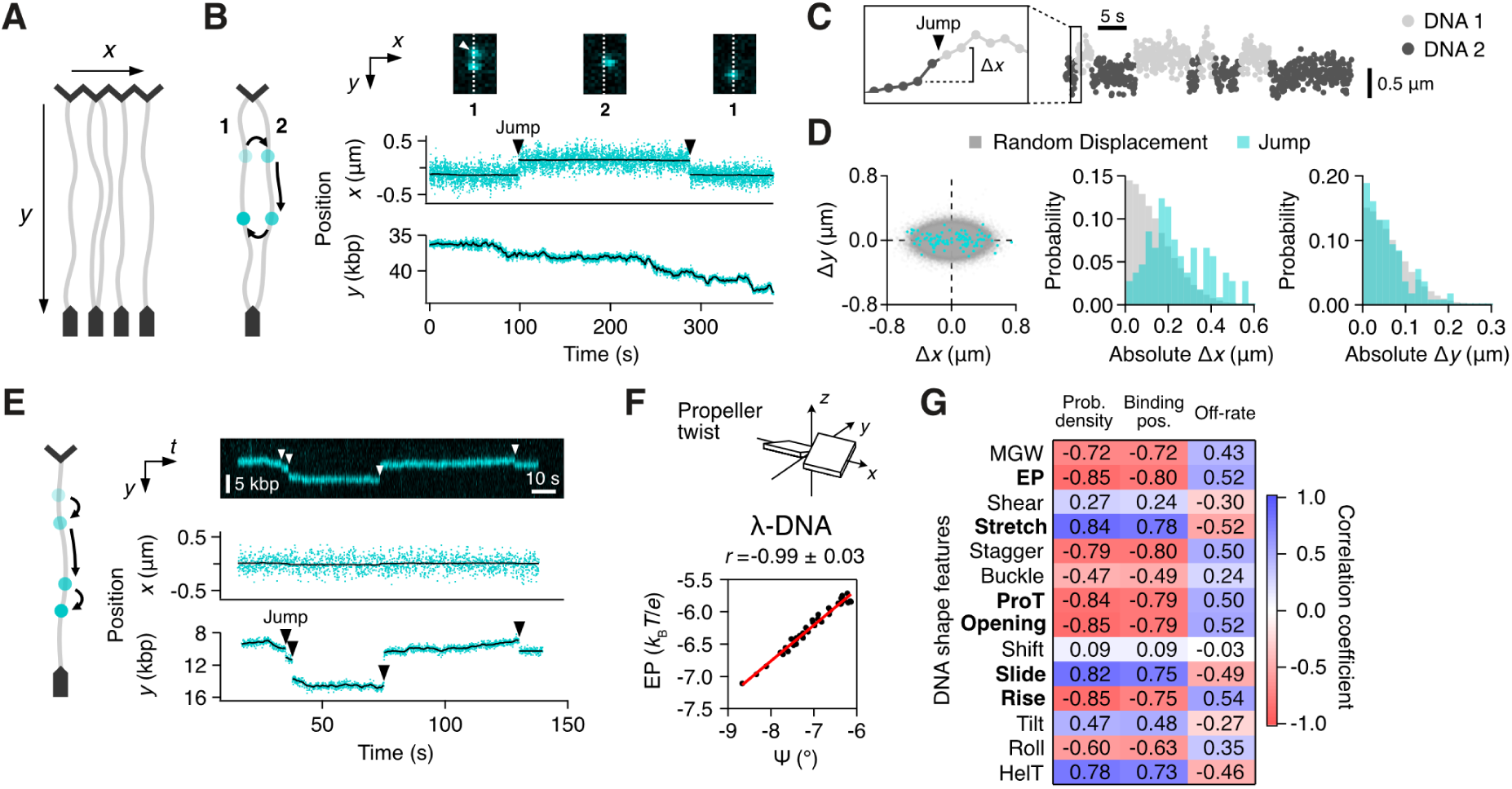
Intersegmental transfers and jumps of INO80 complexes. **(A)** Coordinate system of particle tracking on DNA curtains.**(B)** Example of two intersegmental transfer events by a single INO80 complex. Left: Schematic of the intersegmental transfer events from DNA (1) to (2) to (1). Right top: TIRF microscopy images of the complex before and after each transfer event (striped line represents position of DNA (1). Right bottom: *x*- and *y*-position of the complex over time. **(C)** Analysis of intersegmental transfers. Approximate transfer timepoints were identified manually with assistance of a HMM. To increase accuracy the displacements before and after the identified timepoints were included for analysis and compared to random three-frame displacements. **(D)** Left: Three-frame displacements in *x* and corresponding displacements in *y* for identified transfers (cyan) and random displacements (grey). Middle: Histogram of absolute *x*-displacements. Right: Histogram of absolute y-displacements. *n*_Jump_ = 105. **(E)** Example of four longdistance jumps of a single INO80 complex along the same DNA strand. White triangles indicate jumps. Same graphs as in (B). **(F)** Top: Scheme showing propeller twisting of a DNA basepair. Bottom: Correlation of average EP and propeller twist of the *λ*-DNA used in experiments (1 kbp window). **(G)** Pearson’s correlation coefficients for correlation of different DNA shape features with DNA binding parameters of INO80 along *λ*-DNA shown in fig. 2G.

**FIG. S3.**
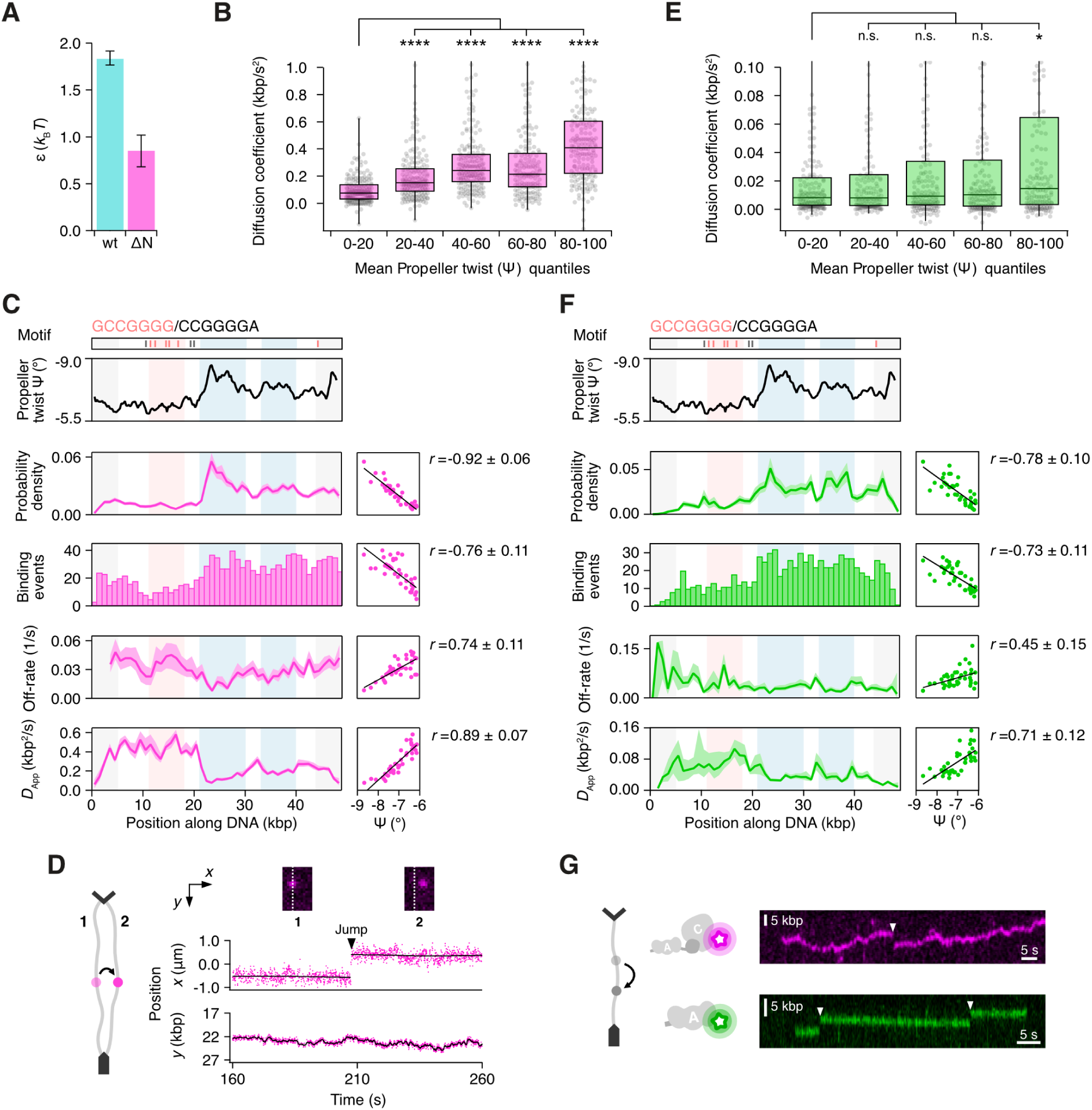
The A-module is a DNA shape sensor. **(A)** Comparison of free-energy roughness for *Sc.* wildtype and *Ct.* ΔN-mutant of INO80. Error bars indicate uncertainty from the range of estimated hydrodynamic radii *R* for both complexes. **(B)** Diffusing *Ct.* ΔN-INO80 complexes were sorted into quintiles depending on the mean propeller twist they encountered during diffusion along *λ*-DNA. Significance was determined using a Mann-Whitney U test. *n* = 877. **(C)** Nhp10-Motifs, average propeller twist (1 kb sliding window) and different binding parameters of *Ct.* ΔN-INO80 along *λ*-DNA. Right boxes display correlation of binding parameters with average propeller twist and Pearson’s correlation coefficient. Shaded areas show chrome barriers (gray), Nhp10 motifs (red) and areas of high negative propeller twist (blue). Grey shaded areas were not included in correlation analysis. *n* = 1180. **(D)** As fig. S2B, but for *Ct.* ΔN-INO80. **(E)** As fig. S3B, but for the A-module. *n* = 783. **(F)** As fig. S3C, but for the A-module. *n* = 851. **(G)** Example of long-distance jumps of single mutant complexes along the same DNA strand for *Ct.* ΔN-INO80 and A-module. White triangles indicate jumps.

**FIG. S4.**
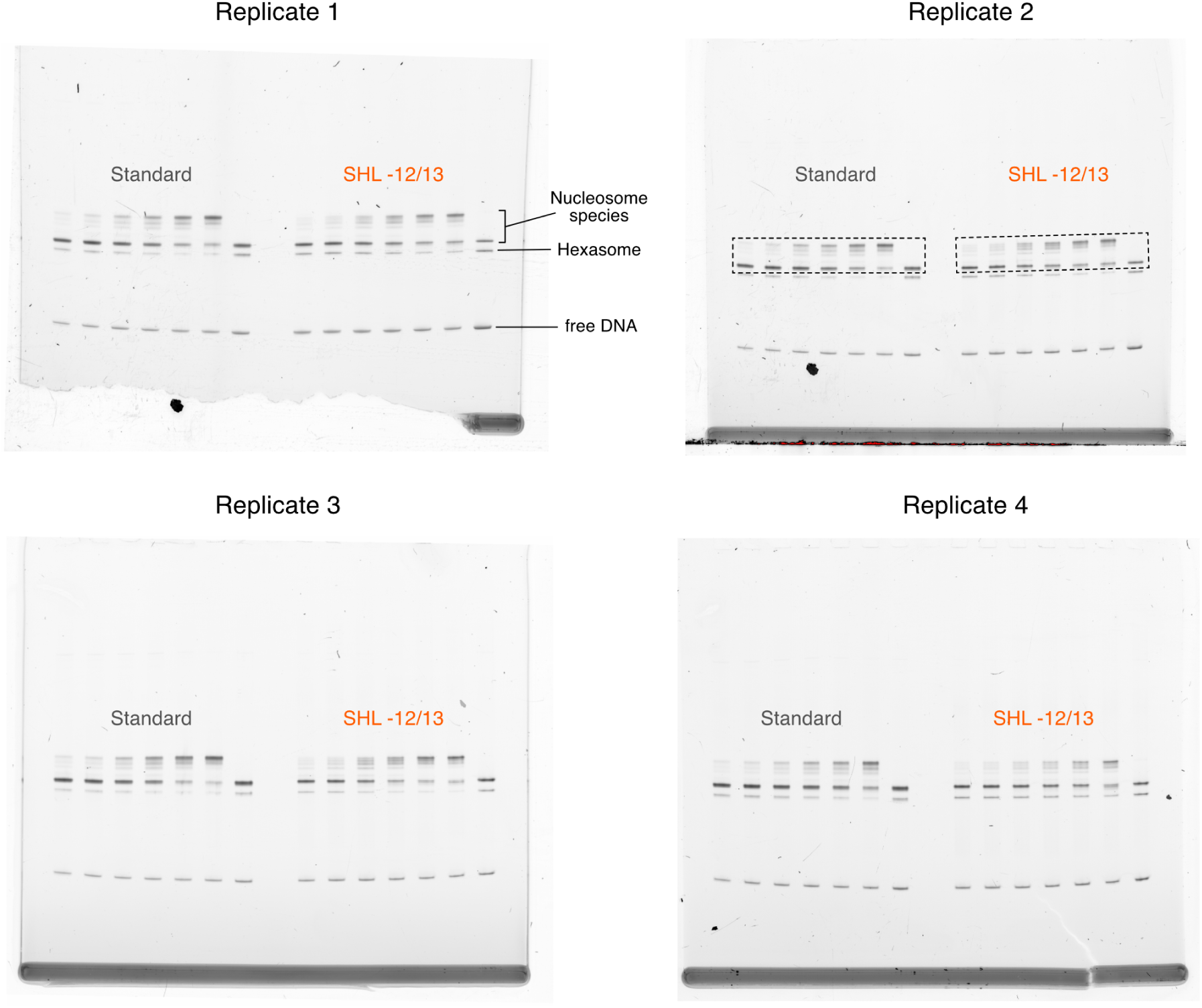
Effect of DNA shape sensing on nucleosome remodeling. Complete gel pictures of INO80 sliding assays. Bands shown in fig. 4B are highlighted by dashed box.

**FIG. S5.**
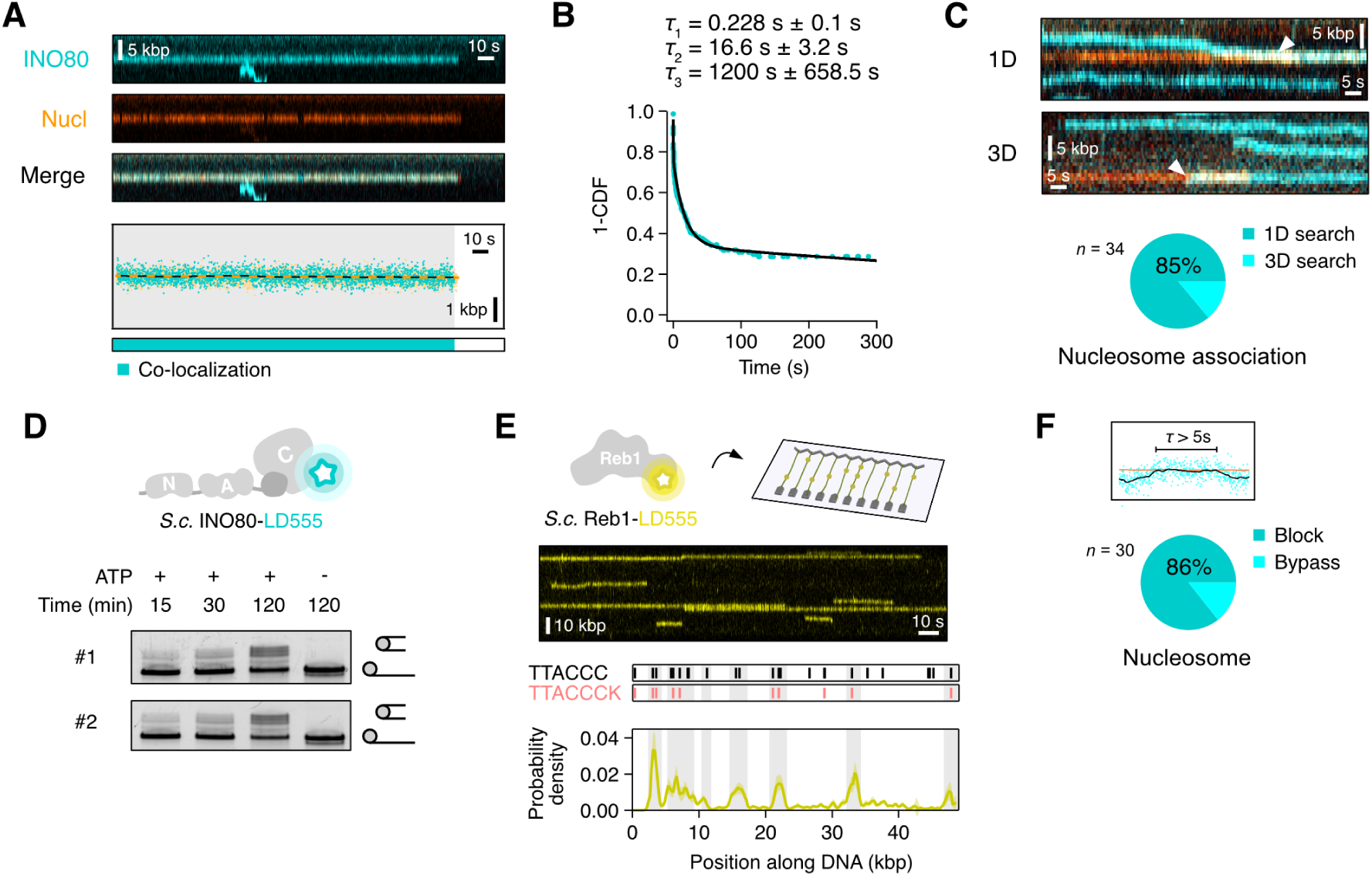
INO80 interactions with nucleosomes and Reb1. **(A)** Top: Kymogram of an INO80 complex (cyan) colocalizing with a nucleosome (orange). Bottom: Corresponding time traces with co-localizations indicated by shaded areas. **(B)** Top: Lifetimes deducted from best fit for Kaplan-Meier curve. Survival plot of INO80-nucleosome co-localization times. Black line represents best fit. **(C)** Sliding assays with INO80-LD555 and 0N80 *H.sapiens* nucleosomes. Two representative replicates shown. **(D)** Top: Schematic of Reb1 and DNA curtains assay. Bottom: Kymogram of Reb1 binding to *λ*-DNA. **(E)** Reb1-binding motifs (stronger binding motifs in red) and probability density of Reb1 along *λ*-DNA. Grey shading indicates peaks of Reb1 binding that correspond with Reb1-binding motifs. *n* = 724. **(F)** Pie chart indicating fraction of INO80-nucleosome co-localization events with a length of over 5 s that result in a block.

